# The *Ruminococcus bromii* amylosome protein Sas6 binds single and double helical α-glucan structures in starch

**DOI:** 10.1101/2022.11.20.514607

**Authors:** Amanda L. Photenhauer, Filipe M. Cerqueira, Rosendo Villafuerte-Vega, Krista M. Armbruster, Filip Mareček, Tiantian Chen, Zdzislaw Wawrzak, Jesse B. Hopkins, Craig W. Vander Kooi, Štefan Janeček, Brandon T. Ruotolo, Nicole M. Koropatkin

## Abstract

Resistant starch is a prebiotic with breakdown by gut bacteria requiring the action of specialized amylases and starch-binding proteins. The human gut symbiont *Ruminococcus bromii* expresses granular starch-binding protein Sas6 (Starch Adherence System member 6) that consists of two starch-specific carbohydrate binding modules from family 26 (RbCBM26) and family 74 (RbCBM74). Here we present the crystal structures of Sas6 and *Rb*CBM74 with a double helical dimer of maltodecaose bound along an extended surface groove. Binding data combined with native mass spectrometry suggest that RbCBM26 binds short maltooligosaccharides while RbCBM74 can bind single and double helical α-glucans. Our results support a model by which RbCBM74 and RbCBM26 bind neighboring α-glucan chains at the granule surface. CBM74s are conserved among starch granule-degrading bacteria and our work provides molecular insight into how this structure is accommodated by select gut species.

## Introduction

The gut microbiota, the consortium of microbes that resides in the human gastrointestinal tract, influences many aspects of host physiology including digestive health [1]. The composition of the gut microbiota is modulated by the human diet [2–4]. After host nutrient absorption in the small intestine, indigestible dietary fiber transits the large intestine and becomes food for gut microbes [3]. Bacterial fermentation of dietary carbohydrates produces beneficial short chain fatty acids including butyrate, a primary carbon source for colonocytes that also has systemic anti-inflammatory and anti-tumorigenic properties [3, 5].

Resistant starch is a prebiotic fiber that tends to increase butyrate in the large intestine [6]. Starch is a glucose polymer composed of branched, soluble amylopectin and coiled insoluble amylose [7, 8]. Breakdown of starch starts with human salivary and pancreatic amylases which release maltooligosaccharides for absorption in the small intestine [9]. However, a portion of starch is indigestible by human amylases and is termed resistant starch (RS) [9]. Raw, uncooked starch granules are resistant to digestion in the upper gastrointestinal tract due to the tight packing of constituent amylose and amylopectin into semi-crystalline, insoluble granules [7]. This type of resistant starch, called RS2, becomes food for gut bacteria that can adhere to and deconstruct granules, releasing glucose and maltooligosaccharides that cross-feed other organisms [9].

Human gut bacteria that degrade RS2 *in vitro* include *Bifidobacterium adolescentis* and *Ruminococcus bromii* [10–14]. *R. bromii* is a Gram-positive anaerobe that increases in relative abundance in the gut upon host consumption of resistant potato or corn starch [10, 15, 16]. *R. bromii* is a keystone species for RS2 degradation because it cross-feeds butyrate-producing bacteria [10]. *R. bromii* synthesizes multi-protein starch-degrading complexes called amylosomes via protein-protein interactions between dockerin and complementary cohesin domains [17–19]. As many as 32 *R. bromii* proteins have predicted cohesin or dockerin domains including amylases, pullulanases, starch-binding proteins, and proteins of unknown function [17, 20]. Many have carbohydrate-binding modules (CBMs) that presumably aid in binding starch and tether the bacteria to its food source [21].

CBMs are classified by amino acid sequence into numbered families and include members that bind only soluble starch and some that also bind granular starch [21, 22]. One such family is CBM74 which was discovered as a discrete domain (*Ma*CBM74) of a multimodular amylase from the potato starch-degrading bacterium, *Microbacterium aurum* [22]. *Ma*CBM74 binds amylose and amylopectin as well as raw wheat, corn, and potato starch granules [22]. The CBM74 family is unique as it is ∼300 amino acids, two to three times larger than most starch-binding CBMs [21]. CBM74 domains are typically found in multimodular enzymes that include a glycoside hydrolase family 13 (GH13) domain for hydrolyzing starch and are flanked by a starch-binding CBM from family 25 or 26 (CBM25 or CBM26) [21, 22]. Most CBM74 family members are encoded by gut microbes and 70% are found in Bifidobacteria [22]. The genomes of *R. bromii* and *B. adolescentis* each encode one putative CBM74-containing protein. The prevalence of CBM74 domains encoded within the genomes of RS2-degrading bacteria, and its increased representation in metagenomic and metatranscriptomic analyses from host diet studies, suggest a role for this module in RS2 recognition in the distal gut [23–25].

The *R. bromii* starch adherence system protein 6 (Sas6) is a secreted protein of 734 amino acids that contains both a CBM26 and CBM74 followed by a C-terminal dockerin type 1 domain [26, 27]. Here we present the biochemical characterization and crystal structure of Sas6, providing the first view of the CBM74 domain and its juxtaposition with the CBM26 domain. The co-crystal structure of *Rb*CBM74 with a double helical dimer of maltodecaose, which mimics the architecture of double helical amylopectin in starch granules, revealed recognition via an elongated groove spanning the domain. *Rb*CBM74 exclusively binds longer maltooligosaccharides (≥ 8 glucose units), and native mass spectrometry suggests that both single and double helical α-glucans are recognized, providing flexible recognition of amylose and amylopectin. Our biochemical data demonstrate that CBM26 and CBM74 recognize different α-glucan moieties within starch granules leading to overall enhanced granule binding.

## Results

### Modular Architecture of Sas6

Sas6 consists of five discrete domains: an N-terminal CBM26 (*Rb*CBM26), a CBM74 domain (*Rb*CBM74) flanked by Bacterial Immunoglobulin-like (BIg) domains, and a C-terminal dockerin type I (**Fig. 1A**) [27]. Sas6 is encoded at the WP_015523730 locus (formerly RBR_14490 or Doc6, UniProt: A0A2N0UYM2) and includes a Gram-positive signal peptide (residues 1-30) that presumably targets the protein for secretion. *Rb*CBM74 spans residues 242-572 based on an alignment with annotated CBM74 domains [22]. We used InterProScan to annotate the remaining sequence which added the Bacterial Immunoglobulin-like (BIg, Pfam 02368) domain A (BIgA), but did not predict BIgB, which we identified via structure determination [28].

**Figure 1:**
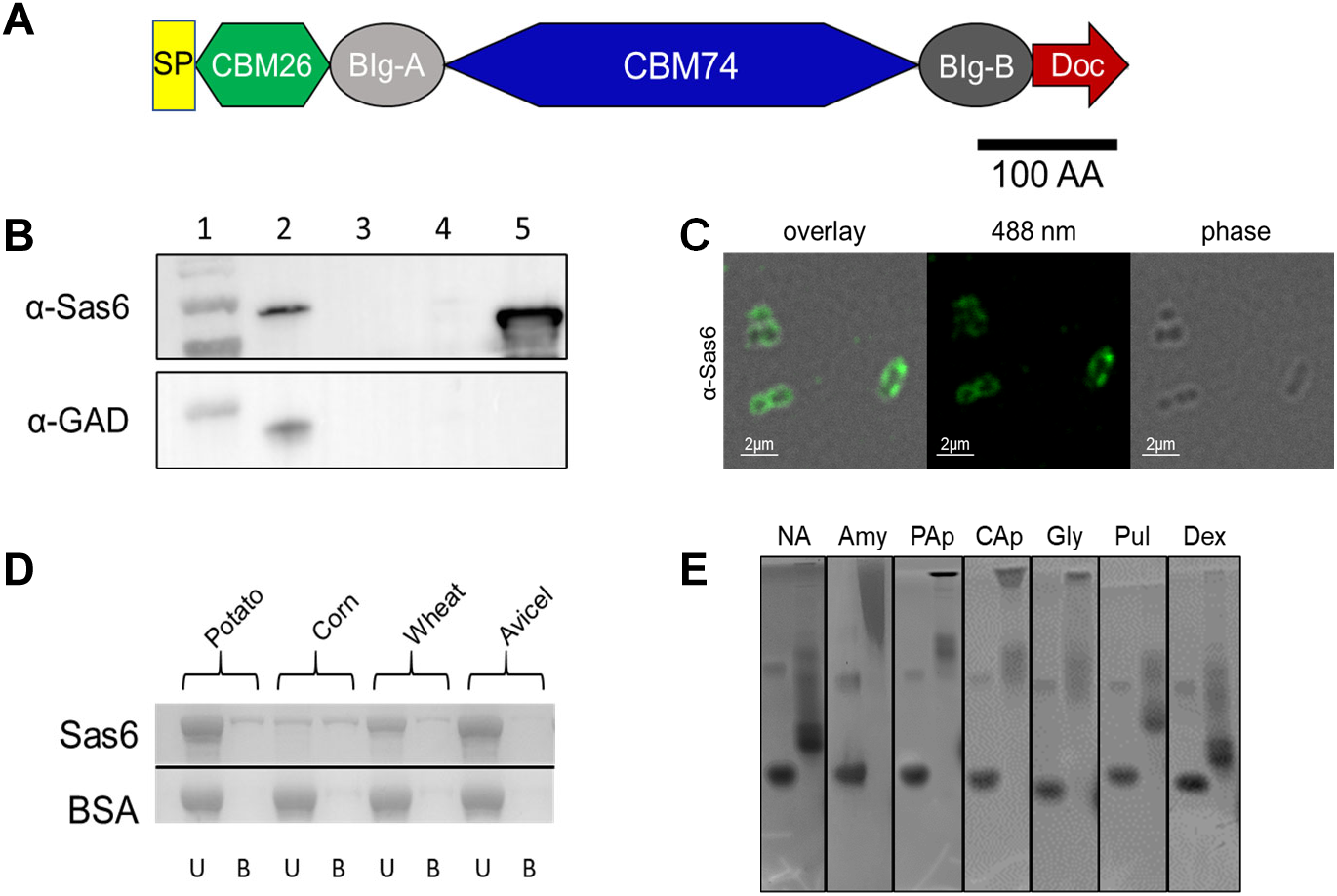
R*u*minococcus *bromii* Sas6 is a starch-binding protein that contains two carbohydrate-binding modules. **A.** Domain architecture of Sas6 annotated according to the Carbohydrate Active Enzyme database (www.cazy.org) and the crystal structure. SP = Signal Peptide, CBM26 = Carbohydrate Binding Module family 26, BIg = Bacterial Immunoglobulin, CBM74 = Carbohydrate Binding Module family 74, Doc = Dockerin. **B.** *Top:* Western blot with anti-Sas6 antibody showing localization of Sas6 in the cell fraction. *Bottom*: Parallel western blot with custom rabbit antiserum against glutamic acid decarboxylase to control for cell lysis. Lane 1: ladder, 2: *R. bromii* cell lysate, 3: cell-free culture supernatant, 4: TCA precipitated cell-free culture supernatant, 5: recombinant Sas6T, truncated version of Sas6 lacking the C-terminal dockerin. **C.** α-Sas6 immunofluorescent staining of fixed *R. bromii* cells grown in potato amylopectin. **D.** SDS-PAGE gel from Sas6 adsorption to potato, corn, and wheat starch, and Avicel (cellulose) control. U=unbound protein, B=bound protein. **E.** Affinity PAGE with 0.1% of the indicated polysaccharide incorporated into the gel matrix. For each, left lane is bovine serum albumin, right lane is Sas6T. NA=native gel, Amy=potato amylose, PAp=Potato Amylopectin, CAp=corn amylopectin, Gly=Glycogen, Pul=Pullulan, Dex=Dextran.

### Sas6 Cell Localization

Though Sas6 has a signal peptide it is unknown whether it is a constituent of a cell-bound amylosome, or part of a freely secreted complex [20]. *R. bromii* synthesizes five scaffoldin (Sca) proteins that have cohesins for amylosome assembly; Sca2 and Sca5 are cell-bound and Sca1, Sca3, and Sca4 are freely secreted [20]. The cognate cohesin for the Sas6 dockerin is unknown. Sas6 is detected in the cell-free supernatant of *R. bromii* cultures in stationary phase but also elutes from the surface of exponentially growing cells with EDTA which disrupts the calcium-dependent cohesin-dockerin interaction [17, 29]. To determine the localization of Sas6, we grew cells to mid-log phase on potato amylopectin and performed a Western Blot with custom antibodies against recombinant Sas6 (**Fig. 1B**). Sas6 was detected in the cell fraction and not the cell-free culture supernatant (**Fig. 1B**), and was visualized on the cell surface via immunofluorescence (**Fig 1C**). Therefore, we conclude that Sas6 is a component of a cell-surface amylosome in actively growing cells. It is possible that Sas6 localization is dependent upon growth phase, as are cellulosome components in some organisms, explaining its previous detection in culture supernatant [17]. Alternatively, *R. bromii*, like some cellulosome-producing bacteria, may release cell-surface amylosomes in stationary phase [30].

### Sas6 Starch Binding

CBM26 and CBM74 are putative raw starch-binding families [22, 31]. Plant sources of granular starch differ greatly in granule organization, including crystallinity (e.g., packing of the long helical chains), length of α1,4-linked chains, amylose location and organization, water content, and trace elements [7]. We used a truncated construct of Sas6 (residues 31-665) lacking the C-terminal dockerin domain, herein called Sas6T, to test Sas6 binding to starch polysaccharides. Sas6T binds potato, corn, and wheat starch granules, with the highest fraction of protein bound to corn starch, and no non-specific binding to Avicel (crystalline cellulose) (**Fig. 1D**). Of note, corn starch has a smaller granule size and therefore a larger surface area to mass ratio [8]. We tested Sas6T binding to amylopectin and amylose, as well as glycogen and pullulan via affinity PAGE. Glycogen is similar to amylopectin with more frequent α1,6 branching (every 6-15 residues for liver glycogen compared to 15-25 residues for amylopectin) [32, 33]. Pullulan is a fungal α-glucan composed of repeating α1,6-linked maltotriose units [34]. Sas6T binds amylose, amylopectin (potato and corn), and glycogen but has less affinity for pullulan suggesting a preference for longer α1,4-linked regions within the polysaccharide (**Fig. 1E**). Sas6T does not bind dextran, a bacterially derived exopolysaccharide of α1,6-linked glucose [35], demonstrating its specificity for starch.

### Structure of Sas6

The structure of Sas6T with α-cyclodextrin (ACX), was determined via single-wavelength anomalous dispersion of intrinsic sulfur-containing residues to a resolution of 1.6Å (*R*work=16.8%, *R*free=21.2%) (**Table 1**). The final model contained two molecules of Sas6T in the asymmetric unit, with four Ca^2+^ per chain and one molecule of ACX bound at the *Rb*CBM26 domain. The Sas6T structure determined with ACX was used to phase a dataset from unliganded crystals (2.2Å, *R*work=19.7%, *R*free=25.5%) (**Table 1**). The overall crystal structure of Sas6T is compact, with *Rb*CBM26, BIgA and BIgB forming an arc over *Rb*CBM74 (**Fig. 2A**).

**Figure 2:**
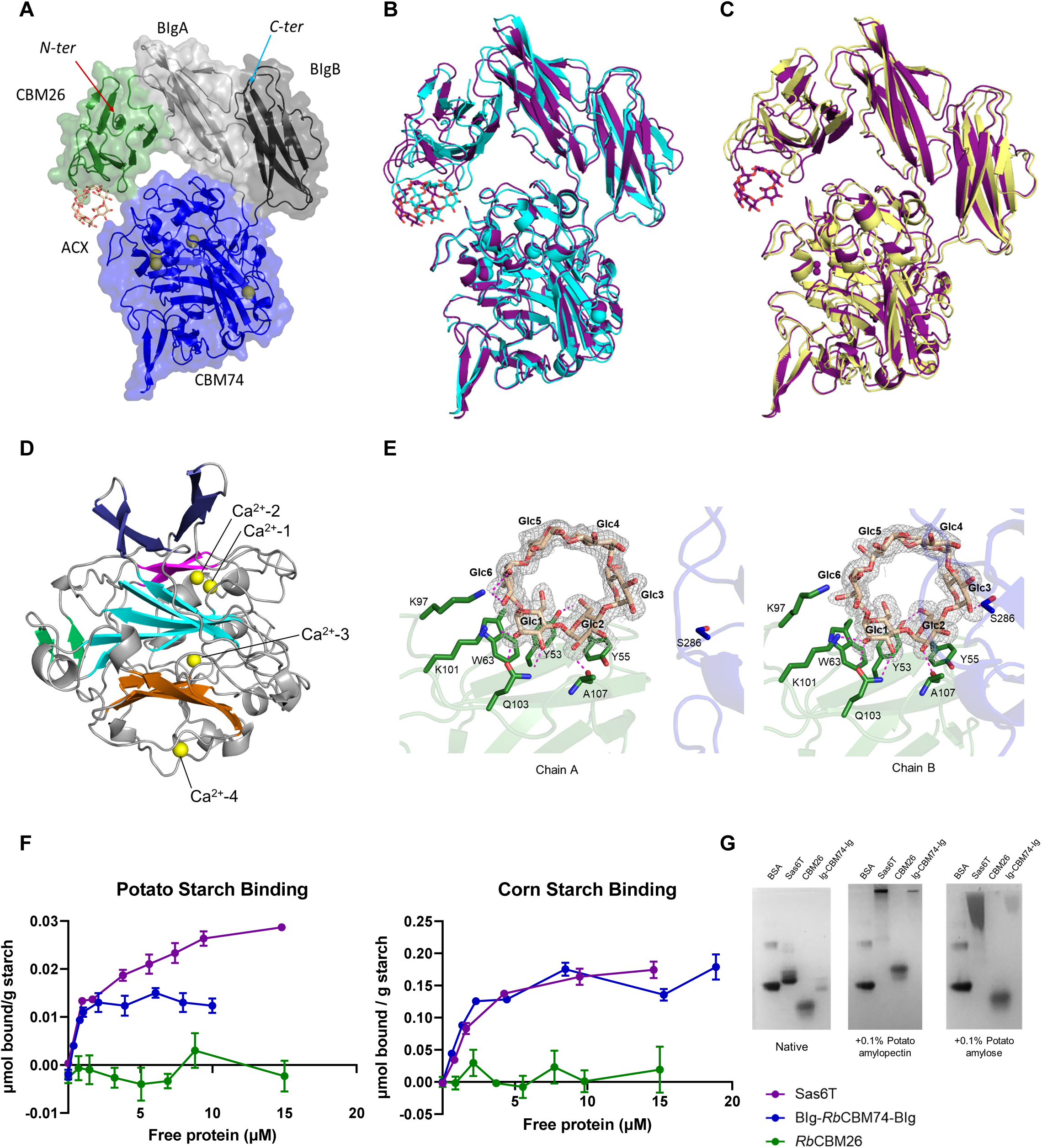
Sas6 is a compact protein with two BIg domains that orient *Rb*CBM26 and *Rb*CBM74. **A.** Semi-transparent surface rendition and cartoon of Sas6T (PDB *7uww*) with *Rb*CBM26 domain in green, BIgA in light grey, *Rb*CBM74 in blue, and BIgB in dark grey. The α-cyclodextrin (ACX) bound to *Rb*CBM26 is shown in wheat sticks and Ca^2+^ atoms are shown as yellow spheres. **B.** Overlay of Chain A (purple) and Chain B (cyan) within the asymmetric unit of *7uww* showing variation in the position of ACX relative to *Rb*CBM74. **C.** Overlay of Chain A of *7uww* (purple) and SAXS-derived MultiFoXS model (yellow). **D.** Side view of *Rb*CBM74 with the central β-sandwich sheets in orange and cyan. A third β-sheet is shown in magenta and the protruding pairs of β-strands and in dark blue. β-strands connecting the beginning and end of the *Rb*CBM74 domain are colored green. Ca^2+^ atoms are shown as yellow spheres. **E.** ACX bound at *Rb*CBM26 (green) in chain A (left) and chain B (right), demonstrating minor conformational flexibility that places S286 from *Rb*CBM74 (blue) within the binding site. Side chains involved in ligand binding are shown as green sticks with a hydrogen bond cutoff of 3.2Å. ACX is displayed as wheat sticks. Omit map is contoured to 2.0σ and carved within 1.6Å of ACX ligand. **F.** *Rb*CBM74 drives binding to granular potato and corn starch. Binding to granular starch was determined by isotherm depletion. The μmoles of protein bound per gram of starch was plotted against [free protein] to determine dissociation constants (Kd) and binding maxima (Bmax) using a one-site specific binding model in GraphPad prism. **G.** Affinity PAGE of Sas6T or individual domains, *Rb*CBM26 and BIg-*Rb*CBM74-BIg, with 0.1% polysaccharide. BSA= bovine serum albumin.

**Table 1:** X-ray Data Collection and Refinement Statistics

| Table 1: X-ray Data Collection and Refinement Statistics |  |  |  |
| --- | --- | --- | --- |
| Construct | Sas6T + α-cyclodextrin | Sas6T unliganded | Blg-RbCBM74-Blg + G10 |
| PDB Accession | 7UWW | 7UWU | 7UWV |
| Wavelength | 0.979 | 0.979 | 0.979 |
| Resolution range | 35 - 1.61 (1.67 - 1.61) | 44.77 - 2.19 (2.27 - 2.19) | 62.48 - 1.70 (1.76 - 1.70) |
| Space group | P 21 21 21 | P 21 21 21 | P 21 21 2 |
| Unit cell | 69.5 82.5 213.5 90 90 90 | 69.2 82.4 213.3 90 90 90 | 69.7 160.1 67.8 90 90 90 |
| Total reflections | 1690626 (29309) | 1044914 (104226) | 512772 (51008) |
| Unique reflections | 122358 (2727) | 63042 (6261) | 84071 (8261) |
| Multiplicity | 13.8 (10.7) | 16.6 (16.9) | 6.1 (6.2) |
| Completeness (%) | 76.8 (24.8) | 99.34 (99.97) | 99.9 (100.0) |
| Mean I/sigma(I) | 16.0 (1.1) | 22.66 (14.34) | 17.2 (2.2) |
| R-merge | 0.092 (1.799) | 0.0956 (0.192) | 0.052 (0.75) |
| R-meas | 0.095 (1.889) | 0.0987 (0.1979) | 0.057 (0.81) |
| R-pim | 0.025 (0.563) | 0.0243 (0.04781) | 0.023 (0.33) |
| CC1/2 | 0.999 (0.549) | 0.998 (0.993) | 0.999 (0.822) |
| Reflections used in refinement | 122315 (3895) | 63244 (6262) | 84061 (8260) |
| Reflections used for R-free | 6093 (201) | 3200 (309) | 4056 (427) |
| R-work | 0.168 (0.238) | 0.197 (0.276) | 0.179 (0.279) |
| R-free | 0.212 (0.281) | 0.255 (0.348) | 0.199 (0.281) |
| Number of non-hydrogen atoms | 11425 | 10523 | 4954 |
| macromolecules | 9721 | 9527 | 4043 |
| ligands | 253 | 38 | 237 |
| solvent | 1451 | 964 | 674 |
| Protein residues | 1294 | 1246 | 531 |
| RMS(bonds) | 0.013 | 0.001 | 0.013 |
| RMS(angles) | 1.4 | 0.4 | 1.7 |
| Ramachandran favored (%) | 96.3 | 96.1 | 97 |
| Ramachandran allowed (%) | 3.6 | 3.9 | 3 |
| Ramachandran outliers (%) | 0.08 | 0 | 0 |
| Rotamer outliers (%) | 0 | 1.6 | 0.5 |
| Clashscore | 3.26 | 4.98 | 0.24 |
| Average B-factor | 22.2 | 25.6 | 33.2 |
| macromolecules | 20.7 | 25.4 | 31.8 |
| ligands | 35.8 | 29.9 | 31.5 |
| solvent | 30 | 27.8 | 41.8 |

*Rb*CBM26, *Rb*CBM74, and the dockerin domain are separated by BIgA (light grey) and BIgB (dark grey), respectively (**Fig. 2A, Extended Data Fig 1A**). Ig-like or fibronectin-III domains act as spacers in multi-modular glycoside hydrolases including GH13s that target starch [36]. BIgA and BIgB interact via hydrogen bonding with 354Å of buried surface area [37] (**Extended Data Fig 1B**). This interaction may help stabilize or orient the CBM74 domain or the BIgs may act as a hinge between the CBMs. The two chains in the asymmetric unit exhibit some flexibility resulting in different positioning between the *Rb*CBM26 binding site and the *Rb*CBM74 domain (**Fig. 2B**).

### Small Angle X-Ray Scattering

To better connect how our crystal structures correlate with conformational flexibility in solution, we used size-exclusion chromatography coupled with small angle x-ray scattering (SEC-SAXS) on Sas6T (**Table 1**). The elution separated out several peaks, including a single strong peak for that was well separated and monodisperse as indicated by the constant radius of gyration (Rg) across the eluted peak (**Extended Data Fig 2A**). The Guinier fit of a subtracted scattering profile created from that peak gave Rg and I(0) values of 29.44 ± 0.04Å and 0.04 ± 3.65 x 10^-5^ and the fit and normalized fit residuals confirmed this peak was monodisperse (**Extended Data Fig 2B**). The molecular weight of Sas6T from the SAXS data was calculated to be 61.0 kDa (theoretical 68.9 kDa) indicating it is primarily monomeric in solution [38]. The Dmax from the P(r) function for Sas6T is 90Å. The overall shape of the P(r) function for Sas6T, calculated by indirect Fourier transform (IFT) using GNOM, has a relatively Gaussian shape that is characteristic of a globular compact particle with the main peak at r = ∼30 Å (**Extended Data Fig 2C**) [39]. There is a small peak at r = 55Å which suggests there are two structurally separate motifs, possibly *Rb*CBM26 and *Rb*CBM74. The dimensionless Kratky plot maxima for Sas6T are typical for a rigid globular protein (**Extended Data Fig 2D**). The small plateau in the mid to high q region, around qRg = 5 in the dimensionless Kratky plot indicates some extension or disorder in the system. These results suggest the presence of two separate modules with flexibility between them, likely corresponding to the two CBMs.

We tested whether the crystal structure matched the solution data by fitting the crystal structure to the SAXS data using FoXS [40]. The fit had a χ^2^= 2.46 and showed systematic deviations in the normalized fit residual (**Extended Data Fig 2E**). This highlights that there are significant differences between the lowest energy conformation of Sas6T in the crystal structure and the structure of Sas6T in solution. We then used MultiFoXS with our high-resolution structure of Sas6T to account for the flexibility, assigning the linkers between the domains (residues 130-137 and 572-583) as flexible [40]. MultiFoXS gave a best fit with a 1-state solution with a χ^2^=and calculated Rg of 29.2Å which corroborates the Guinier Rg calculation (**Extended Data Fig 2F**). An alignment of Chain A of the crystal structure and MultiFoXS model had a RMSD of 1.2Å over 347 pruned atom pairs (**Fig. 2C**). The MultiFoXS model shows a slightly more extended model for Sas6T in comparison to the crystal structure demonstrating that Sas6T has some flexibility in solution yet remains compact.

*Structure of* Rb*CBM74 – Rb*CBM74 (357 residues) has 21 β-strands and 13 short α-helices with a core β-sandwich fold of two sheets with five antiparallel β-strands (**Fig. 2D, Extended Data Fig 3A**). A third short β-sheet forms a convex face and two pairs of β-strands (residues 356-369 and 412-423) protrude from the region between the β-sandwich and the third β-sheet. In this structure, two short β-strands lie at the entrance and exit of the CBM74 domain, marking the domain boundaries (**Extended Data Fig 3B**).

**Figure 3:**
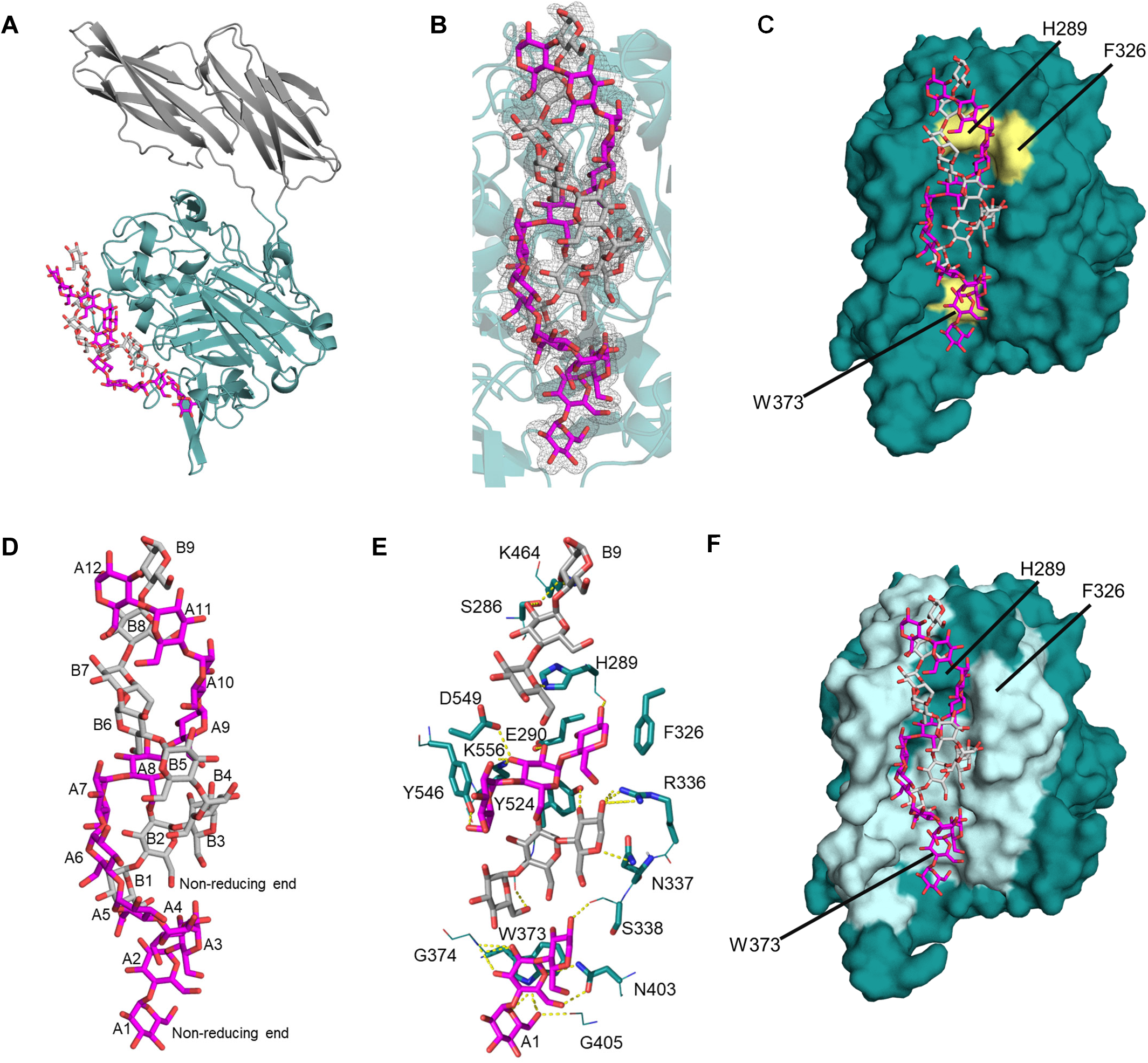
R*b*CBM74 has an extended groove that accommodates starch double helices. **A.** The BIg-*Rb*CBM74-BIg (PDB *7uwv*) starch-binding site is an extended groove that spans nearly the length of the domain. A cartoon representation of BIgA in light grey, CBM74 in teal, and BIgB in dark grey with two chains of maltodecaose (G10) wrapped around one another shown in magenta and grey sticks. **B.** *Rb*CBM74 is co-crystallized with G10 in a double helical conformation. Electron density for G10 demonstrated by an omit map contoured to 2.0σ and carved to 1.6Å with one chain of modeled Glc in magenta and the other in grey. **C.** *Rb*CBM74 has an inset binding groove that accommodates the width of the starch double helix with aromatic CH-π stacking provided by W373, F326, and H289. A surface representation of CBM74 (teal) with aromatic residues colored in yellow and G10 represented by magenta and grey sticks. **D.** Double helical G10 structure with Glc residues labeled from non-reducing to reducing ends. One chain of G10 (A1-12) shown in magenta and the other in grey (B1-9) sticks. **E.** Corresponding hydrogen-bonding network (3.2Å cutoff) between *Rb*CBM74 and G10. Side chains involved in hydrogen bonding are shown in teal sticks with nitrogens indicated in blue and oxygens in red. Hydrogen bonds are indicated by yellow dashed lines and G10 residues directly involved in binding are shown in magenta (G10 molecule A) and grey (G10 molecule B) sticks. **F.** Surface representation of *Rb*CBM74 with peptides protected from deuterium exchange in the presence of G10 colored in light cyan as determined by hydrogen-deuterium exchange mass spectrometry.

A DALI search revealed that the central fold of *Rb*CBM74 most closely resembles CBM9 from *Thermotoga maritima* Xylanase10A (PDB ID: 1I82-A, Z-score: 9.8, RMSD: 3.2Å, identity: 17%) [41, 42] (**Extended Data Fig 3C**). *Tm*CBM9 binds glucose, cellobiose, cello- and xylo-oligomers at the reducing ends, and amorphous and crystalline cellulose [42]. *Tm*CBM9 (189 residues) is larger than most CBMs which range from 80-120 amino acids [42]. Despite the conserved core β-sandwich, *Rb*CBM74 displays several extra loops and β-strands. The ligand binding site of *Tm*CBM9 is formed by two Trp residues that create an aromatic clamp around cellobiose. *Rb*CBM74 W373 is conserved with one of these Trps and lies within an extended, shallow channel partially covered by residues 374-384 that form a flexible loop only resolved in one monomer (**Extended Data Fig 3D**).

There are three putative structural Ca^2+^ in the *Tm*CBM9 structure and four cations in RbCBM74, one of which aligns with a Ca^2+^ in *Tm*CBM9 (**Extended Data Fig 3E**). We modeled these cations as Ca^2+^ based upon coordination geometry and atomic distances (**Extended Data Fig 3F**) [43, 44]. Ca^2+^-1 and Ca^2+^-2 are separated by 3.8Å and share three coordinating residues but only Ca^2+^-2 is surface exposed. Ca^2+^-3 is abutted by the loop connecting β-strands 2 and 3 and Ca^2+^-4 is at the center of a loop formed by residues 256-264 and conserved with *Tm*CBM9. Like *Tm*CBM9, the Ca^2+^ ions in the *Rb*CBM74 structure may be important for structural stability [45].

### Molecular Basis of RbCBM26 Binding

The N-terminal *Rb*CBM26 displays a β-sandwich consistent with other members of the CBM26 family [21]. In both chains of the asymmetric unit, CH/π stacking with ACX is provided by W63 and Y55 with hydrogen bonding mediated by Y53, K101, Q103, and the peptidic oxygen of A107 (**Fig. 2E**). In chain A only, K97 provides hydrogen bonding with O3 of Glc6. In chain B, ACX lies 3.2Å from S286 of the CBM74 domain and hydrogen bonds with O2 and O3 of Glc3. In contrast, S286 is 9.5Å from ACX in chain A. The top structural homologs of *Rb*CBM26 from DALI are the CBM25 from *Bacillus halodurans* C-125 (*Bh*CBM26) from α-amylase G-6 (PDB ID: 2C3V-A, Z-score: 12.4, RMSD 1.9Å, identity: 16%) and CBM26 (*Bh*CBM26) from the same enzyme (PDB ID: 6B3P-B, Z-score: 12.1, RMSD 1.9Å, identity: 20%) [41, 46]. Another top DALI result is *Er*CBM26b of Amy13K from *Eubacterium rectale* (PDB ID 2C3H-B, Z-score: 10.8, RMSD 1.7Å, identity: 19%). In all three CBM26 structures, the structure and aromatic platforms for ligand recognition are conserved (**Extended Data Fig 4AB**). *Rb*CBM26, in contrast to *Er*CBM26 and *Bh*CBM26, has a longer loop containing K97 and K101 that provide additional hydrogen bonding with ACX. Unlike *Bh*CBM26, RbCBM26 does not undergo a conformational change upon ligand binding (**Extended Data Fig 4C**) [31]. A sequence alignment with CBM26 members *Bh*CBM26, *Er*CBM26 and the *Lactobacillus amylovorus* α– amylase CBM26 (*La*CBM26), demonstrates conservation of the aromatic platform but more variation in the hydrogen-bonding network (**Extended Data Fig 4A**). Sas6 W63 corresponds to *La*CBM26 W32 that, when mutated, results in complete loss of binding [47]. The *R. bromii* protein Sas20 has a CBM26-like domain that shares 26% sequence identity with *Rb*CBM26, yet *Rb*CBM26 shares more structural similarity with *Bh*CBM26 and *Er*CBM26 [29].

**Figure 4:**
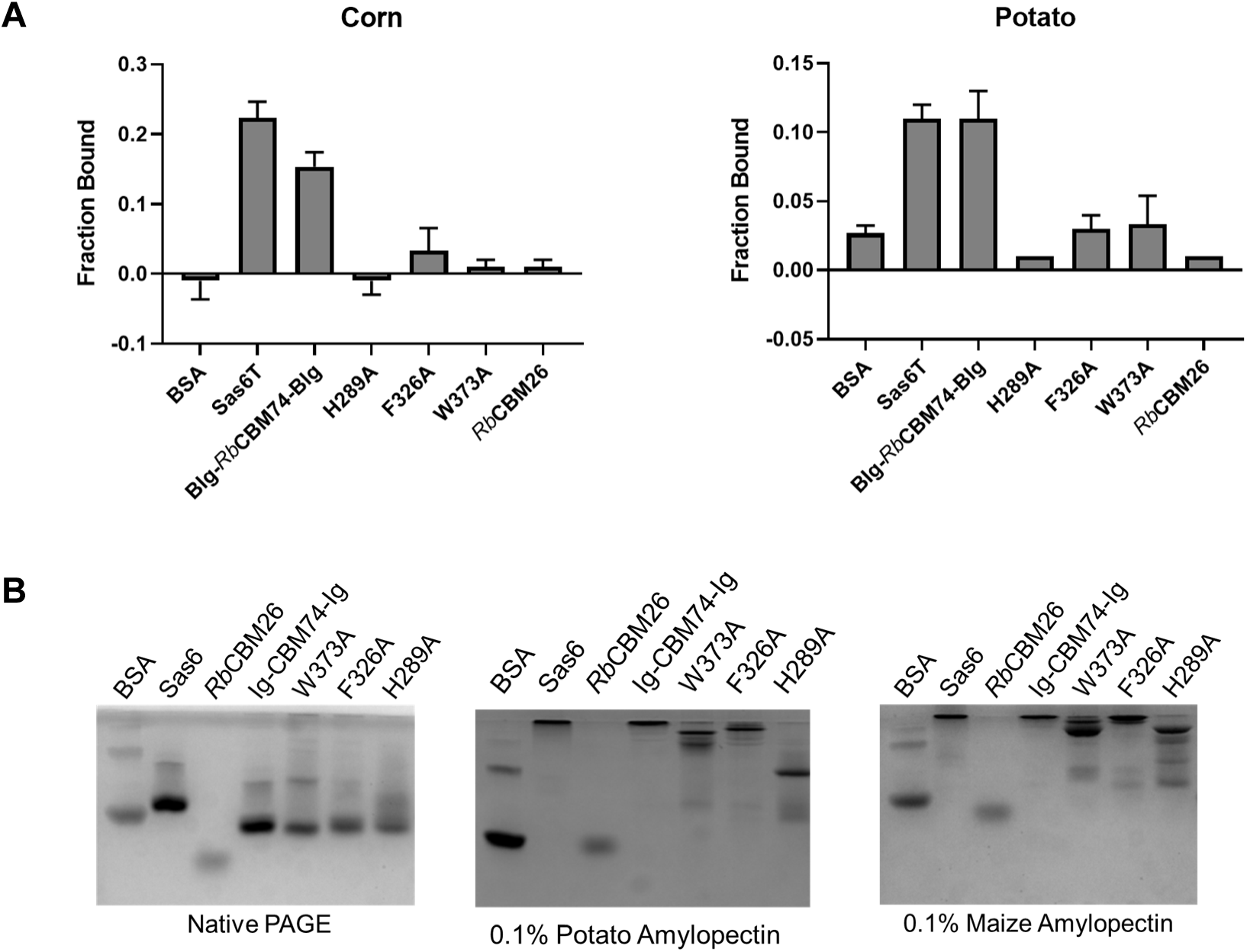
W373A, F326A, and H289A mediate starch binding by *Rb*CBM74. **A.** Binding to insoluble starch is eliminated or greatly reduced when W373, H289 or F326 is mutated. The amount of protein bound to starch granules was determined by quantitation of protein remaining in solution after binding (n = 3). **B.** Mutation of aromatic residues decreases but does not eliminate binding to amylopectin. Affinity PAGE with 0.1% potato amylopectin or maize amylopectin added to the gel matrix. Binding is indicated by reduced migration through the gel.

### Binding Mechanism of Sas6

We expressed the individual Sas6 CBMs and included the BIgA/B domains with the CBM74 (BIg-*Rb*CBM74-BIg, residues 134-665) to enhance solubility. Sas6T and BIg-*Rb*CBM74-BIg bound granular corn and potato starch, but *Rb*CBM26 did not bind either insoluble starch at detectable levels (**Fig. 2F**). Sas6T binds to more of the corn starch granule, (Kd = 2.8µM ± 0.4, Bmax = 0.21µmol/g ± 0.01) but has a modestly higher affinity for potato starch (Kd = 1.9µM ± 0.3, Bmax = 0.030µmol/g ± 0.001), which might be a function of the smaller granule size and larger surface to mass ratio for corn starch. Exclusion of the *Rb*CBM26 in the BIg-*Rb*CBM74-BIg construct led to slightly better binding to corn starch (Kd = 1.5µM ± 0.3, Bmax = 0.18µmol/g ± 0.008) and modestly higher affinity but less overall binding to potato starch (Kd = 0.51µM ± 0.13, Bmax = 0.015µmol/g ± 0.001). The saturation curve for BIg-*Rb*CBM74-BIg closely resembles that of Sas6T and there is minimal binding by *Rb*CBM26, suggesting that *Rb*CBM74 drives insoluble starch binding.

The molecular patterns on the surface of starch granules differs between plant sources and remains an active area of research [48–51]. The “hairy billiard ball model” to describe starch granules postulates that the granule surface has block-like clusters of amylopectin chains with hair-like extensions of amylose penetrating through the amylopectin [50]. Sas6T and BIg-*Rb*CBM74-BIg bind amylose and amylopectin whereas *Rb*CBM26 only binds to amylopectin with apparently low affinity based upon the relatively small change in migration (**Fig. 2G**). This suggests that *Rb*CBM74 drives binding of Sas6 to the long, tightly packed helices of amylose at the surface of the starch granule.

Using isothermal titration calorimetry (ITC), we found that Sas6T and BIg-*Rb*CBM74-BIg bound amylopectin with sub-micromolar affinity whereas binding was not detectable for *Rb*CBM26 (**Table 2**; **Extended Data Fig 5A**) [52]. Sas6T binds maltotriose (G3), maltoheptaose (G7), maltooctaose (G8) with a *K*d in the hundreds of µM but exhibits a Kd of ∼5µM for maltodecaose (G10) (**Table 2**; **Extended Data Fig 5B**). Interestingly, *Rb*CBM26 binds shorter linear oligosaccharides (G3, G7) and cyclodextrins, while BIg-*Rb*CBM74-BIg had no detectable affinity for these sugars (**Table 2**; **Fig. Extended Data Fig 5C**). None of the constructs bound glucosyl-α1,6-maltotriosyl-α1,6-maltotriose, an oligosaccharide of pullulan, suggesting that the α1,6 linkages are not specifically recognized by either domain. We determined that BIg-*Rb*CBM74-BIg binds exclusively longer α-glucans of at least 8 residues. Notably, α1,4-linked glucose polymers form double helices at 10 glucose units due to internal hydrogen bonding so we hypothesized that *Rb*CBM74 might accommodate starch helices [8].

**Table 2:**
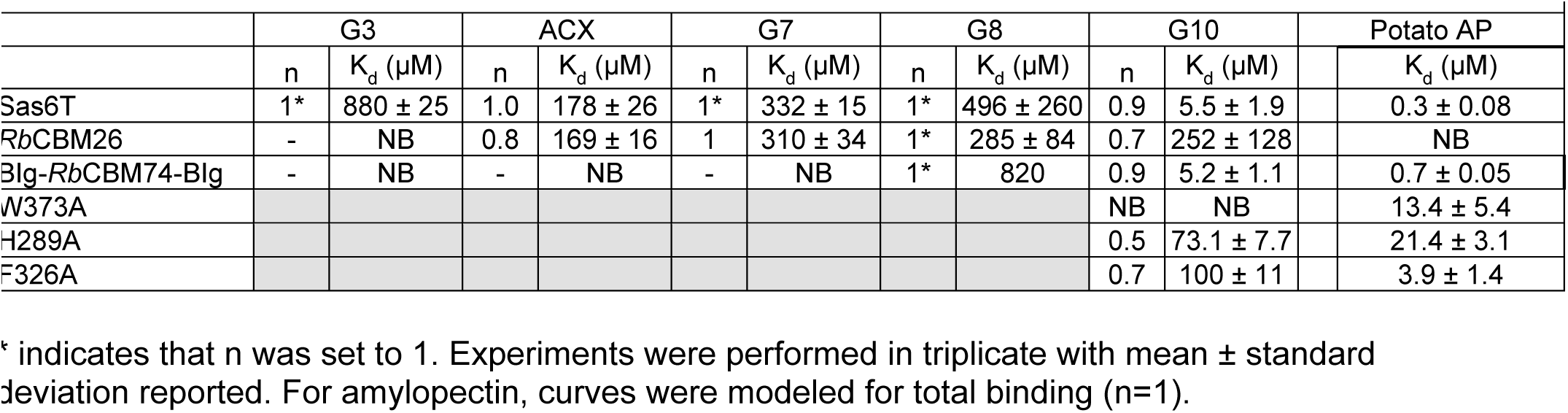
Sas6 and domain binding via Isothermal Titration Calorimetry

**Figure 5:**
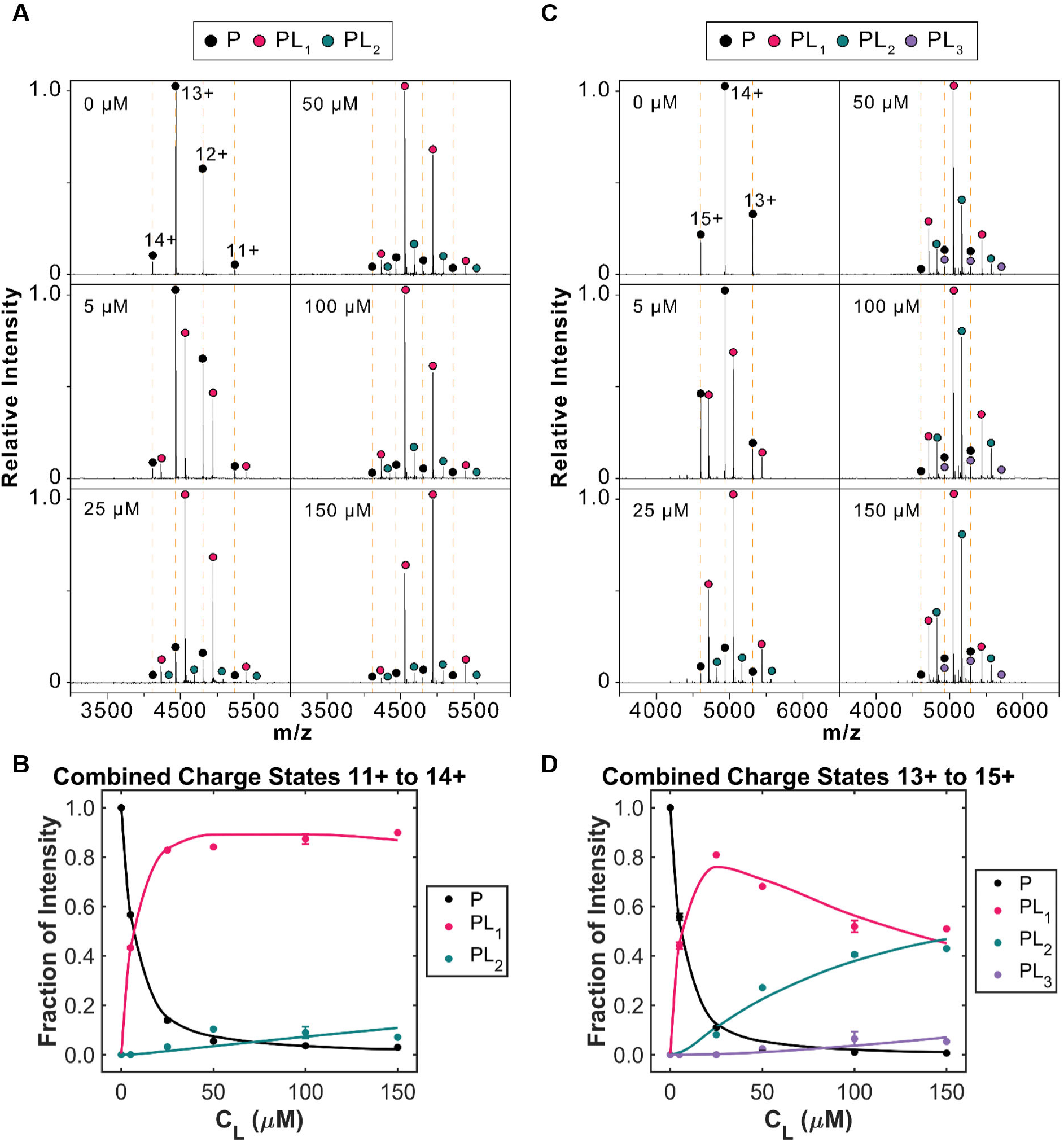
R*b*CBM74 and *Rb*CBM26 bind separate molecules of G10 in solution. **A.** Mass spectra of BIg-*Rb*CBM74-BIg at different ligand concentrations (0 - 150µM) and a fixed protein concentration of 5µM. Charge states for unbound protein are annotated with an orange dashed line. Peaks corresponding to different bound states are observed after each charge state of the unbound protein. Intensities of each species, combined across multiple charge states, were then extracted from the mass spectra and used to calculate the fractional abundance of unbound and bound states at equilibrium (n=3). **B.** Nonlinear least-squares fitting of the titration data for BIg-*Rb*CBM74-BIg. **C.** Mass spectra of Sas6T as described in A. **D.** Nonlinear least-squares fitting of the titration data for Sas6T.

### Molecular Basis of RbCBM74 Binding

We co-crystallized BIg-*Rb*CBM74-BIg with maltodecaose (G10) to 1.70Å resolution (*R*work=17.9%, *R*free=19.9%) (**Fig. 3A**). Remarkably, we observed two molecules of G10 as an extended double helix of ∼42Å along the face of *Rb*CBM74 extending from S286 (reducing ends) to W373 (non-reducing ends). There was strong electron density for 12 glucoses in one molecule, and nine glucoses in the other chain, likely reflecting varied occupancy of the helix along the binding cleft (**Fig. 3B**). H289, F326, and W373 stood out as surface exposed aromatic residues that might be providing CH-π mediated stacking (**Fig. 3C**).

An overlay of the unliganded and G10 bound structures demonstrates little global change in the CBM74 domain upon binding (**Extended Data Fig 6A**), with the exception of G374 to K381. In the unliganded structure this loop occludes surface exposure of W373 and in the G10 bound structure the loop opens to create a continuous binding surface (**Extended Data Fig 6B**). Additionally, Ca^2+^-4 is exchanged for Na^+^, representing flexibility in ion identity at that site (**Extended Data Fig 6C**).

**Figure 6:**
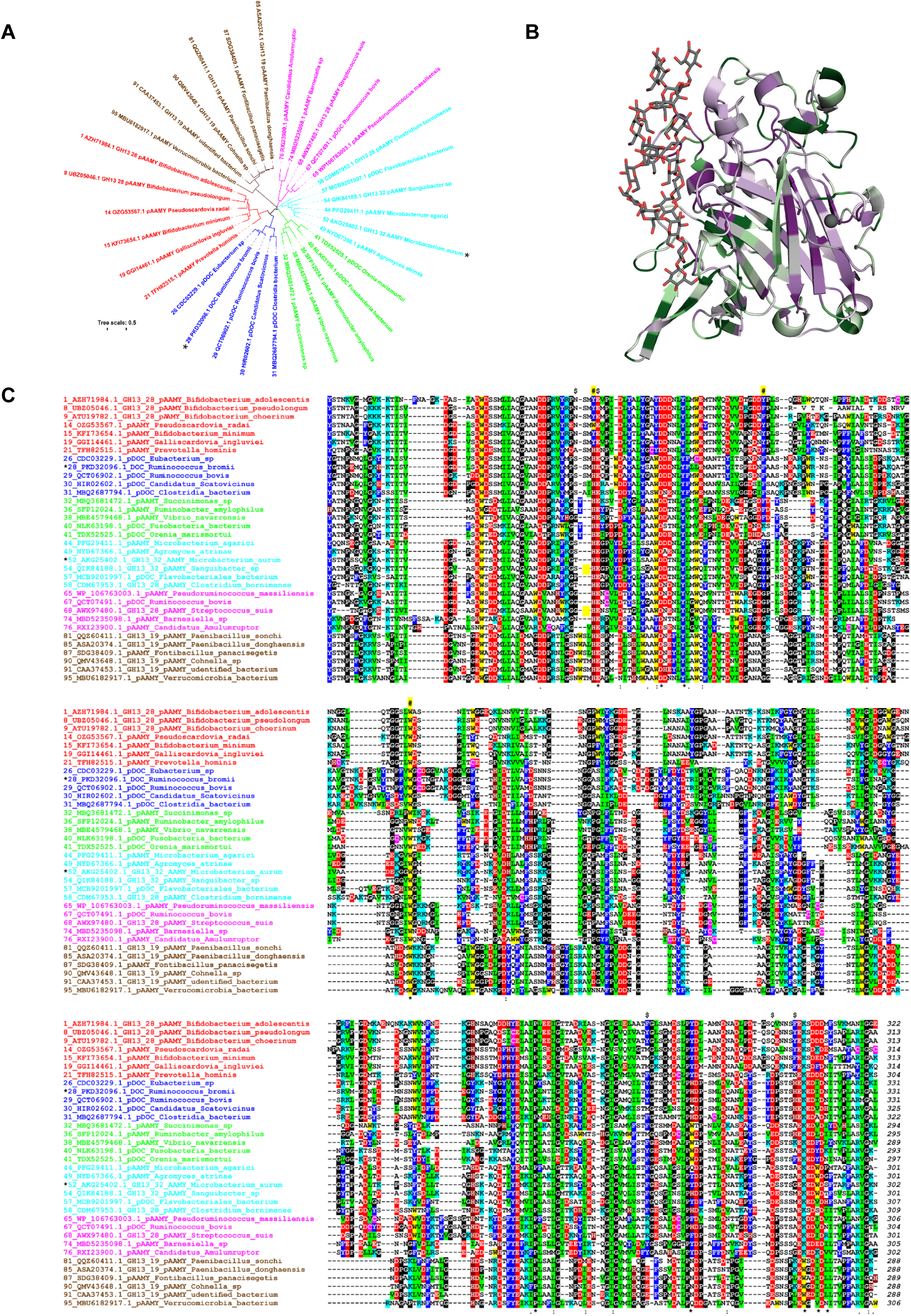
Conservation of binding residues among select CBM74 family members. **A.** Evolutionary tree for the CBM74 family including 33 sequences selected from the entire studied set of 99 CBM74s (Extended Data Table 2). Two experimentally characterized CBM74s are marked by an asterisk: Sas6 from *Ruminococcus bromii* (No. 28, blue cluster) and the subfamily GH13_32 α-amylase from *Microbacterium aurum* (No. 52; cyan cluster). Protein labels include the order number (33 selected from 1-99), GenBank accession number, abbreviation of the source protein/enzyme and organism name. The tree is based on the alignment (shown in C) spanning the complete CBM74 sequences. **B.** Structure of *Rb*CBM74 (PDB *7uwv*) colored by conservation score from least conserved (green) to most conserved (purple) generated using CONSURF. **C.** Sequence alignment of the CBM74 family. The six individual groups distinguished from each other by different colors correspond to six clusters seen in the evolutionary tree (panel A); the sequence order in the alignment reflects their order in the tree in the anticlockwise manner (starting from the first sequence in the red cluster). The residues responsible for stacking interactions and involved in hydrogen bonding with glucose moieties of the bound α-glucan are signified by a hashtag and a dollar sign, respectively, above the alignment. The flexible loop observed in the three-dimensional structure of the *Rb*CBM74 is highlighted by the short yellow strip over the alignment. Identical and similar positions are signified by asterisks and dots/semicolons under the alignment blocks. The color code for the selected residues: W, yellow; F, Y – blue; V, L, I – green; D, E – red; R, K – cyan; H – brown; C – magenta; G, P – black. The alignment of all 99 CBM74 sequences of the present study shown in Extended Data Figure 9B.

Canonical starch-binding domains feature two or three aromatic residues for pi-stacking interactions with the aglycone face of maltooligosaccharides, but *Rb*CBM74 is designed for extensive hydrogen-bonding interactions with longer oligosaccharides and starch [21]. The binding site is continuous and each G10 molecule interacts with protein as a stretch of three Glcs at a time, before the natural helical curvature brings the chain out of the contact with the protein (**Fig. 3D**). For example, at the non-reducing end, Glc 1-3 of G10 chain A (G10A) fit into the ligand-binding groove, while Glcs 4-6 of G10A are solvent exposed and Glc 1-3 of G10 chain B (G10B) then fill the cavity. Along the length of the cavity, from the non-reducing end to the reducing end, Glcs 1-3 and 7-9 of both G10A and G10B alternate to fill this binding site.

The binding cleft features a network of residues that hydrogen bond to the hydroxyl groups of glucose (**Fig. 3E**). At the non-reducing end, Glc A1 hydrogen bonds with the indole nitrogen of W373. Glc A2 stacks with W373 with hydrogen bonding provided by G374 and N403. Glc A3 hydrogen bonds with S338. The other molecule of G10 (B) contacts the next part of the binding groove and is anchored by hydrogen bonding of Glc B3 by R336 and Y524. Where the first molecule turns back into the binding groove, Glc A8 hydrogen bonds with E290, D549, and K556. Glc A9 hydrogen bonds with the backbone of H289 and pi stacks with F326. The H289 side chain hydrogen bonds with Glc B7 and provides aromatic character for pi stacking with Glc B8. Near the region of *Rb*CBM74 that lies adjacent to *Rb*CBM26, K464 and S286 hydrogen bond with Glc B9.

To define the starch-binding properties of *Rb*CBM74 in solution, we employed Hydrogen– Deuterium eXchange Mass Spectrometry (HDX-MS). The conformational dynamics of BIg-*Rb*CBM74-BIg alone and in the presence of G10 were measured over a 4-log timescale (**Extended Data Fig 7AB**). The overall conformational dynamics of the apo protein were consistent with the determined crystal structure, in terms of well-ordered domains and associated loops or flexible regions. The flanking BIg domains showed higher exchange rates than the core CBM74 domain. Intriguingly, the linker regions between domains do not show differentially high dynamic exchange, as would be expected for flexibly tethered independent domains, further supporting the integral nature of BIg-*Rb*CBM74-BIg motif.

The binding of G10 to *Rb*CBM74 was explored by differential protection from exchange in the absence and presence of G10. Significant protection was observed in the presence of G10, while no significant increases in exchange were observed (**Extended Data Fig 7C**). This is consistent with the minimal global conformation changes between the two states of the protein. The protected regions upon G10 binding were highly localized to a single surface binding region (**Fig. 3F**). This protected region constitutes a single extended surface, which directly overlaps with the G10 binding site observed in the co-crystal structure (**Fig. 3EF**). With the exception of the peptide from A314-Y318 (ANTTY), each of the protected peptides identified by HDX-MS contains at least one key binding residue identified from the co-crystal structure (**Fig. 3E**). These data provide a comprehensive picture of the structural dynamics of *Rb*CBM74 binding to long maltooligosaccharides via an extended starch binding cleft.

### RbCBM74 Mutational Studies

Because most CBM binding is mediated by aromatics, we hypothesized that mutation of W373, F326, or H289 to Ala would dramatically decrease or eliminate binding. We tested maximum binding of each of the aromatic mutants to insoluble corn (1%) and potato starch (5%). The W373A and H289A constructs lost the ability to bind to insoluble corn starch while binding of the F326A construct was greatly reduced (**Fig. 4A**). This trend was somewhat different for potato starch, in which a lower percentage of H289A bound compared to the F326A and W373A mutants. By affinity PAGE, neither the W373A nor the F326A mutant lost appreciable binding to amylopectin while the H289A mutant had a modest decrease in binding to potato amylopectin (**Fig. 4B**). When we quantified binding via ITC, W373A lost all binding for G10 while H289A and F326A had a ∼10-20-fold decrease in affinity (**Table 2, Extended Data Fig 8A**). On potato amylopectin, F326A had a 10-fold reduction in affinity while H289A and W373A exhibited a ∼20-fold reduction (**Extended Data Fig 8B**). That single mutations do not eliminate binding is perhaps not surprising given the extensive binding platform. Moreover, the enhanced affinity of these mutants to amylopectin over G10 further suggests that productive interactions with the protein extend beyond a 10-glucose unit footprint. Indeed, the somewhat staggered double helical G10 bound in our crystal structure suggests that at least 12 glucose units contribute to binding (**Fig. 3D**).

### Native mass spectrometry

ITC revealed a binding stoichiometry of 1:1 between BIg-*Rb*CBM74-BIg and G10, while the co-crystal structure demonstrates that two molecules of G10 are accommodated. To better determine the stoichiometry of this binding event, we employed native mass spectrometry in the presence of varying concentrations of G10 (**Fig. 5A**). Each observed state differed by ∼1639 Da, which agrees with the theoretical mass of G10 (**Extended Data Table 1A**). To obtain binding affinities, we summed the peak intensities of all abundant charge states in our mass spectra and analyzed these intensity values as described previously [53] (**Extended Data Table 1B**). The *K*d for BIg-*Rb*CBM74-BIg was determined to be 3.8 ± 0.5µM, which agrees with our ITC data. As the concentration of ligand is increased, ligand molecules can bind nonspecifically during the nESI process, generating artifactual peaks in the mass spectra corresponding to a two ligand-bound complex (**Fig. 5A**). We speculate that in excess concentrations of G10, the molecules can form double helices that are accommodated by the *Rb*CBM74 binding site but that the single molecule binding event represents the most common binding conformation (**Fig. 5B**).

Because Sas6 encodes both a CBM74 and a CBM26, and this co-occurrence is evolutionarily well-conserved, we speculated that *Rb*CBM26 and *Rb*CBM74 could either bind separate G10 molecules or that one ligand could span the region between the two CBM binding sites [22]. We used native mass spectrometry to determine the number of G10 molecules bound to Sas6T, which includes both CBMs. The binding state distribution was markedly different when *Rb*CBM26 was included (**Fig. 5C**). At low G10 concentrations, there is a mix of unliganded, 1-bound, and 2-bound states unlike BIg-*Rb*CBM74-BIg alone (**Fig. 5D**). As G10 increases, the apo and 1-bound states decrease as the 2-bound fraction increases. For Sas6T, *K*d values for 1:1 and 1:2 protein:ligand complexes were calculated to be 3.4 ± 0.5 µM and 165.6 ± 38.8 µM, respectively, and are in reasonable agreement with ITC data (**Extended Data Table 1B**). Together these results suggest that *Rb*CBM26 and *Rb*CBM74 each bind one molecule of G10 independently in solution. In the context of a starch granule, this supports a model whereby each CBM of Sas6 binds adjacent α-glucan chains rather than attaching to the same chain in a continuous manner. Moreover, the propensity for BIg-*Rb*CBM74-BIg to bind a single helix of G10 at low ligand concentrations, as also observed with ITC, suggests that this binding platform prefers single helical α-glucans such as amylose, though it can also tolerate double helical stretches of amylopectin.

### CBM74 Conservation

To visualize conserved features of CBM74 domains, two alignments and a corresponding evolutionary tree were prepared. The first alignment includes all 99 CBM74 sequences (**Extended Data Fig 9A,B**; **Extended Data Table 2**) while the second was simplified for viewing and includes 33 representative CBM74 sequences (**Fig. 6**). Both alignments reveal that the CBM74s fall into 6 distinct clades (**Fig. 6A**; **Extended Data Fig 9A**). *Rb*CBM74 (No. 28) is in a distinct cluster of proteins (blue) that invariably include a dockerin domain as part of the full-length protein. However, there are other CBM74 domains originating from dockerin-containing proteins found in three more groups (green, cyan, and magenta). The prototypical CBM74 of the subfamily GH13_32 α-amylase from *Microbacterium aurum* (No. 52) bins into a clade (cyan) with its GH13_32 counterpart from *Sanguibacter* sp. (No. 54) and the CBM74-containing α-amylase from *Clostridium bornimense* (No. 58). A similar GH13_28 α-amylase from *Streptococcus suis* (No. 68) is in the adjacent cluster (magenta) very close to the CBM74 domains from two other hypothetical dockerin-containing proteins from *Ruminococcus bovis* (No. 67) and Ruminococcaceae bacterium (No. 70). Most CBM74 domains appended to α-amylases from the subfamily GH13_28, predominantly from Bifidobacteria, group together in a separate cluster (red). Finally, the sixth cluster (walnut) covers CBM74 domains found in GH13_19 α-amylases. In total, CBM74 domains occur in α-amylases from several subfamilies or non-catalytic dockerin-containing proteins and are widely represented among Bifidobacteria.

We mapped the conservation of all 99 CBM74 family members onto our structure using CONSURF [54–56] (**Fig 6B**). While the central β-sandwich, ion-coordination sphere, and ligand binding site are highly conserved, the flexible loop in *Rb*CBM74 (residues 373-384) occluding the binding site is more variable (**Fig. 6C**; **Extended Data Fig. 9B**). In all but the three or four most closely related CBM74 sequences – covering only the two genera of *Ruminococcus* and Eubacterium – this loop is short or not present, though how this feature correlates with binding is unknown.

Most of the key aromatic residues that mediate starch-binding in *Rb*CBM74 are highly conserved (**Fig. 6C**). W373 from *Rb*CBM74 is 100% conserved among all 99 identified CBM74 family members (**Extended Data Fig 9B**), while H289 is shared with 78 sequences or substituted with a Tyr (18/99) in Bifidobacteria and *Candidatus* s*catavimonas* (No. 25) and a Trp (3/99) in *Pseudoscardovia* species. F326 is perhaps the most variable, sharing sequence identity or similarity with 3 of the 6 clades (F-19/99, Y-43/99), while the other clades feature a glycine or alanine in this position (36/99). The binding site also features an elaborate network of residues that provide hydrogen bonding with the ligand. The residues at the center of the cleft including K556 (80/99), D549 (63/99), and E290 (99/99) exhibit the highest conservation (**Fig. 6C**; **Extended Data Fig 9B**). The hydrogen bonding residues at the ends of the cleft are more varied, including S286 (22/99) which interacts with the *Rb*CBM26 ligand. Intriguingly, in a large proportion of the sequences there is an aromatic residue at the site of K556 (W-19/99) and Y524 (Y-12/99, F-45/99) that could provide pi stacking in those CBM74s. This moderate variability in the composition of the putative binding site may suggest that CBM74 family members have different affinities for starch.

## Discussion

CBMs are distinct protein domains that assist with substrate breakdown by specifically binding polysaccharide targets. These domains are especially important for binding to insoluble substrates like crystalline cellulose and semi-crystalline starch granules. The CBM74 family binds insoluble starch and its constituents, amylose and amylopectin. CBM74 domains are frequently (81/99 sequences) encoded adjacent to another starch-binding CBM family, either a CBM25 or CBM26 [22]. Sas6 includes both a CBM26 and a CBM74 domain that have different affinities for maltooligosaccharides but work together to bind granular starch. *Rb*CBM26 has a canonical binding platform that accommodates motifs found in linear and circular maltooligosaccharides. In contrast, *Rb*CBM74 has an extended ligand binding groove that requires at least 8 glucose residues and accommodates the single helices of amylose and the double helices found in amylopectin. Because it is on the cell surface, the CBM74 domain of Sas6 may target *R. bromii* to the crystalline regions of starch granules that are not easily accessible to human or other bacterial amylases.

Sas6 is a putative *R. bromii* amylosome component and likely cooperates with amylases and pullulanases via the interaction of its dockerin domain with a cohesin from a scaffoldin protein [20]. Because Sas6 is found on the cell surface, it could bind cell anchored scaffoldins Sca2 or Sca5, associate with Sca1/Amy4, or bind the cell surface in a dockerin-independent mechanism [20]. Breakdown of starch by *R. bromii* relies on the coordinated effort of approximately 40 distinct proteins, of which Sas6 may play an integral part by specifically targeting the helical regions of starch [20].

Unlike *R. bromii*, resistant starch-utilizing Bifidobacteria encode CBM74-containing multimodular extracellular amylases [9]. A recent study looked at the amylases that were differentially encoded between Bifidobacterial strains that could bind and degrade starch granules and those that could not [57]. <u>R</u>esistant <u>S</u>tarch <u>D</u>egrading enzyme 3 (RSD3) was differentially encoded in the resistant starch-binding strains. It contains a CBM74 domain and has high activity on high amylose corn starch. RSD3 has an N-terminal GH13 domain followed by CBM74, CBM26, and CBM25 domains. The CBM74-CBM26 motif is present in RSD3 so the structural and functional insights we have gleaned from Sas6 may suggest how these CBMs structurally assist the enzyme with granular starch hydrolysis.

Although starch is a polymer composed solely of glucose, there is massive variation in granule structure [7, 8]. This is a function of primary structure (i.e. α1,4 or α1,6 linkages), secondary structure (single or double helices) and tertiary structure (helical packing and amylose content), making granules an exquisitely complex substrate [58]. This complexity is unlocked by only a few specialized gut bacteria, making granular starch a targeted prebiotic [9, 15, 16]. CBM74 domains might serve as a molecular marker for the ability to break down resistant starch in metagenomic samples [22]. Furthermore, CBM74 domains might make attractive additions to engineered enzymes for enhanced starch degradation on the industrial scale, or as an adjunct to starch prebiotics. The structural and functional picture of *Rb*CBM74 here will accelerate the targeted use of this domain for various health and industrial applications.

## METHODS

### Recombinant Protein Cloning and Expression

We used a previously described cloning and expression protocol to generate each of the recombinant protein constructs used in this study [59]. Genomic DNA was isolated from *R. bromii* strain L2-63 and the constructs for Sas6 without the signal peptide were amplified using the primers listed in **Table S1** with overhangs complementary to the Expresso T7 Cloning & Expression System N-His pETite vector (Lucigen). The forward primers were engineered to include the 6x His sequence that complemented the vector plus a TEV protease recognition site for later tag removal. PCR was performed with Flash PHUSION polymerase (ThermoFisher). The amplified products and the linearized N-his pETite vector were transformed in HI-Control10G Chemically Competent Cells (Lucigen) and plated on LB plates supplemented with 50 μg/ml kanamycin (Kan). Transformants were screened for the insertion of Sas6 and validated via sequencing. The Sas6-pETite plasmids were transformed into chloramphenicol (Chl)-resistant *E. coli* Rosetta (DE3) pLysS cells and plated on LB plates supplemented with 50μg/ml Kan and 20μg/mL Chl. *E. coli* cells were grown at 37°C to OD600 0.6-0.8 in Terrific Broth supplemented with 50μg/ml Kan and 20μg/ml Chl after which time the temperature was lowered to 20°C and 0.5mM Isopropyl β-d-1-thiogalactopyranoside (IPTG) was added. After 16 hours of growth, 1L of cells was centrifuged, resuspended in 40mL of Buffer A (20mM Tris pH 8.0, 300mM NaCl) and lysed by sonication. Cell lysate was separated from cell debris by centrifugation for 30min at 30,000xg. 3mL of Ni-NTA resin was packed into Econo-Pac Chromatography Columns (BioRad) and equilibrated with Buffer A. Lysate was passed through the packed columns and washed with 70mL of Buffer A. Proteins were eluted from the columns via stepwise increase in Buffer B (20mM Tris pH 8.0, 300mM NaCl, 500mM imidazole). Proteins eluted in 10-25% Buffer B fractions. TEV protease (1mg) was added to each protein to initiate cleavage of the His-tag and the mix was dialyzed overnight using dialysis tubing (SpectraPor) in 1L of storage buffer (20mM HEPES pH 7, 100mM NaCl). The dialyzed protein-TEV mixture was applied to Ni-NTA resin and the flow-through was collected and concentrated using a VivaSpin 20 concentrator (Fisher Scientific).

### Sas6 Immunofluorescence

Custom α-Sas6T antiserum was generated by rabbit immunization with purified recombinant Sas6T protein (Lampire Biological Laboratories). The resulting antiserum was used for western blotting and cell staining. *R. bromii* cells were grown to mid-log phase on RUM media [17] with 0.1% potato amylopectin and 2mL of the cell culture was collected for immunostaining and western blotting. For immunostaining, 1mL of *R. bromii* culture was centrifuged for 1min at 13,000xg and washed 3 times with 1X phosphate buffed saline pH 7.4 (PBS). 2µL of cells were then spread on a glass slide and fixed with 10% formaldehyde in PBS. Slides were washed 3x in PBS to remove fixative but were not permeabilized. Cells were blocked for 30min with 10% goat serum (Jackson ImmunoResearch). α-Sas6T antiserum was diluted 1:1000 in 10% goat serum and applied for 1hr to cells at room temperature. The primary antiserum was removed, and slides were washed 3 x 5min in PBS before the application of 1:500 goat α-rabbit AlexaFluor488 antibody (ThermoFisher) for 30min. Slides were washed 3 x 5min in PBS and preserved with Prolong Gold Antifade reagent and dried overnight before imaging. Cells were imaged at the University of Michigan Microscopy Core on a Leica Stellaris Light Scanning Confocal microscope with a 100X objective.

### Western Blotting

*R. bromii* was grown to mid-log phase overnight on RUM media containing 0.1% potato amylopectin [17]. 1mL of cells was pelleted and washed twice in phosphate buffered saline (PBS) pH 7.4, then resuspended to a final volume of 50µL in 5mM Tris-HCl pH 8.5. The culture supernatant was passed through a 0.2µm filter and 50µL was reserved for analysis. Proteins were precipitated from the remaining supernatant by the addition of ¼ volume of 100% trichloroacetic acid (TCA) and incubated 30 min on ice. The precipitate was collected via centrifugation and washed twice with 200µL cold acetone. The resulting pellet was dried and resuspended in 50µL of 5mM Tris-HCl pH 8.5. Samples were separated by SDS-PAGE on two 10% Tris-glycine gels, then transferred to polyvinylidene difluoride (PVDF) membrane. Blots were blocked in EveryBlot Blocking Buffer (BioRad) for 30min then washed with PBS pH 7.4 + 0.05% Tween 20 (PBST). To detect Sas6, one membrane was incubated with custom rabbit α-Sas6 antiserum (Lampire) diluted 1:500 and the other with custom rabbit α-glutamic acid decarboxylase from *R. bromii* (Lampire) diluted 1:10,000 in PBST + 5% non-fat dry milk (PBST-milk) for 1hr. Blots were washed in PBST and incubated in horse radish peroxidase-conjugated goat α-rabbit antibody (ThermoFisher) diluted 1:5,000 in PBST-milk and the signal was detected by ECL chemiluminescence (ThermoFisher).

### Granular starch binding assays

Granular starch-binding assays were conducted with potato starch (Bob’s Red Mill), corn starch (Sigma), wheat starch (Sigma), or Avicel (Fluka). Prior to use, all polysaccharides were washed 3x with an excess of assay buffer (20mM HEPES pH 7.0, 100mM NaCl) to remove soluble starch and oligosaccharides and prepared as a 50mg/mL slurry. 1mg (corn) or 5mg (potato) of starch slurry was aliquoted into 0.2mL tubes in triplicate, centrifuged at 2,000xg for 2 min and the supernatant was carefully removed. 100μL of protein ranging from 0.5μM-10μM protein was added to each starch and the tubes were agitated by end-over-end rotation at room temperature for 1hr. After centrifugation at 2,000xg for 2min, 20μL of the supernatant was removed for unbound protein concentration determination by absorbance at A280 using a ThermoFisher NanodropOne with three replicate measurements per sample. The remaining 80μL of supernatant was removed and set aside for SDS-PAGE gel analysis. The concentration of unbound protein remaining in the supernatant was used to determine the µmoles of protein bound per gram of starch which was plotted against the concentration of initial (free) protein to generate a binding curve [31]. Overall affinity (Kd) and binding maximum (Bmax) was determined via a one-site binding model (specific binding) using GraphPad Prism version 9.2.0 for Windows (GraphPad Software, San Diego, California USA, www.graphpad.com) [31].

To assess the remaining starch granules for bound protein, the granules were washed three times with an excess of assay buffer by mixing and centrifugation, the final wash supernatant was removed, and 100μL of Laemmli buffer containing 1M urea was added to the starch pellet to denature any bound protein but keep the original volume consistent. To qualitatively determine the amount of unbound and bound protein, 10μL each of the wash supernatant and solubilized pellet fraction were run separately via SDS-PAGE. Bovine serum albumin was used as a negative control and to confirm unbound protein was sufficiently washed from the starch granules.

### Polysaccharide Affinity PAGE

Non-denaturing polyacrylamide gels with and without potato amylopectin (Sigma), corn amylopectin (Sigma), potato amylose (Sigma), bovine liver glycogen (Sigma), pullulan (Sigma), or dextran (Sigma) to a final concentration of 0.1% polysaccharide were cast. All polysaccharides were autoclaved and amylose was solubilized by alkaline solubilization with 1M NaOH and acid neutralization to pH 7 with HCl [60]. Sas6 protein samples were mixed with 6X loading dye lacking SDS. Gels were run concurrently for 4 hours on ice and subsequently stained with Coomassie (0.025% Coomassie blue R350, 10% acetic acid, and 45% methanol). Gels were imaged on a Bio-Rad Gel Doc Go imaging system. The distance between each band and the top of the separating gel were measured using ImageJ [61]. The ratio of the distance migrated by each band was determined to the distance the BSA band traveled. Binding was considered positive if the ratio was less 0.85 as previously described [62].

### Isothermal Titration Calorimetry

All ITC experiments were carried out using a TA Instruments standard volume NanoITC. For each experiment, 1300μL of 25μM protein was added to the sample cell and the reference cell was filled with distilled water. The sample injection syringe was loaded with 250μL of the appropriate ligand concentration (0.5mM - 5mM) to fully saturate the protein by the end of 25 injections of 10μls. Titrations were performed at 25°C with a stirring speed of 250 rpm. The resulting data were modeled using TA Instruments NanoAnalyze software employing the pre-set models for independent binding and blank (constant) to subtract the heat of dilution. For interactions with high affinity (c-value at 25µM protein greater than 5), no alterations were made to the model. If the calculated c value of an interaction fell below 5, the n value was set to 1 as indicated in the figure legend following the guidance for modeling low affinity interactions [63]. For polysaccharide titrations, curves were modeled by varying the substrate concentration until n=1 such that the Kd represents the overall affinity for the construct [52].

### Protein Crystallization

Crystallization conditions for α-cyclodextrin (2mM) bound (pdb 7UWW) and unliganded (pdb 7UWU) crystals of Sas6T were screened via 96-well sparse matrix screen (Peg Ion HT, Hampton Research #HR2-139) in a sitting drop vapor diffusion experiment at room temperature. Screens were set up using an Art Robbins Gryphon robot with 20mg/mL protein in a 3-well tray (Art Robbins #102-0001-13) using protein-to-well solution ratios of 2:1, 1:1, and 1:2. Small crystals were observed in 0.2M Potassium thiocyanate pH 7.0, 20% w/v Polyethylene glycol 3,350 (condition B2) and were further optimized by varying pH, PEG 3350 percentage, and potassium thiocyanate concentration. Crystals were microseeded with a crystal seeding tool (Hampton) in a sitting drop setup of 1.5µL drops with 2:1, 1:1, or 1:2 protein:well solution ratios. The optimal crystallization solution contained 0.3M Potassium thiocyanate pH 7.0, 24% PEG 3350 and 1mM Anderson−Evans polyoxotungstate [TeW6O24]^6−^ (TEW) (Jena Biosciences #X-TEW-5) to improve crystal diffraction. Prior to data collection, crystals were cryoprotected in a mixture of 80% crystallization solution supplemented with 20% ethylene glycol then plunged into liquid nitrogen.

Crystallization conditions for maltodecaose-bound RbCBM74 structure (pdb 7UWV) were generated from the construct lacking the CBM26 domain (BIg-*Rb*CBM74-BIg, residues 134-665) using 96-well sparse matrix screens. A crystalline mass observed in 60% v/v Tacsimate pH 7.0, 0.1 M BIS-TRIS propane pH 7.0 (Hampton Salt-Rx HT-well H12 #HR2-136) was used to microseed an optimized solution containing 30% Tacsimate, 0.1M HEPES pH 7.0 and 2mM maltodecaose (CarboExpert). No additional cryo-protection was required prior to plunge freezing into liquid nitrogen.

### Structure Determination and Refinement

X-ray data were collected at the Life Sciences Collaborative Access Team (LS-CAT) at Argonne National Laboratory’s Advanced Photon Source (APS) in Argonne, IL. Data were processed at APS using autoPROC with XDS for spot finding, indexing, and integration followed by Aimless for scaling and merging [64–66]. Intrinsic sulfur SAD phasing was used to determine the structure of Sas6T/α-cyclodextrin (7UWW) using AutoSol in Phenix [67, 68]. Those coordinates were then used for molecular replacement in Phaser to determine the unliganded Sas6T (7UWU) and BIg-*Rb*CBM74-BIg/G10 (7UWV) structures [69]. All three structures were refined via manual model building in Coot and refinement in Phenix.refine [70, 71]. Metal ion identities were validated using the web-based CheckMyMetal (CMM) tool [72] (https://cmm.minorlab.org/). Carbohydrate models were validated using Privateer [73].

### SEC-SAXS experiment

SAXS was performed at Biophysics Collaborative Access Team (BioCAT, beamline 18ID at APS) with in-line size exclusion chromatography (SEC-SAXS) to separate the sample from aggregates and other contaminants. Sample was loaded onto a Superdex 200 Increase 10/300 GL column (Cytiva), which was run at 0.6ml/min by an AKTA Pure FPLC (GE) and the eluate after it passed through the UV monitor was flown through the SAXS flow cell. The flow cell consists of a 1.0mm ID quartz capillary with ∼20μm walls. A coflowing buffer sheath is used to separate the sample from the capillary walls, helping prevent radiation damage [74]. Scattering intensity was recorded using a Pilatus3 X 1M (Dectris) detector which was placed 3.6m from the sample giving a q-range of 0.003Å^-1^ to 0.35Å^-1^. 0.7 s exposures were acquired every 1s during elution and data was reduced using BioXTAS RAW 2.1.1 [75]. Within RAW, the Volume of Correlation (*VC*), molecular weight, and oligomeric state were determined [76, 77]. Buffer blanks were created by averaging regions flanking the elution peak and subtracted from exposures selected from the elution peak to create the I(q) vs q curves used for subsequent analyses. The molecular weight was calculated by comparison to known structures (Shape&Size) [38]. P(r) function was determined using GNOM [39]. GNOM and Shape&Size are part of the ATSAS package (version 3.0) [78]. High resolution structures were fit to the SAXS data using FoXS and flexibility in the high-resolution structures was modeled against the Multi-FoXS data [40]. **Tables S2A-C** list sample, instrumentation, and software for the SEC-SAXS experiment.

### Hydrogen–Deuterium eXchange Mass Spectrometry (HDX-MS)

HDX-MS experiments were performed using a Synapt G2-SX HDMS system (Waters), similar to previously reported [79]. Deuteration reactions were incubated at 20°C for 15s, 150s, 1500s, and 15,000s in triplicate. 3μL of BIg-*Rb*CBM74-BIg alone or in the presence of G10 were diluted with 57μL of deuterated labeling buffer. Nondeuterated data were acquired by dilution with protonated buffer and fully deuterated data were prepared by dilution in 99% D2O, 1% (v/v) formic acid) for 48h at room temperature. Samples were measured in triplicate using automated handling with a PAL liquid handling system (LEAP), using randomized sequential collection with Chronos.

Following incubation, deuteration was quenched by mixing 50μL of the solution with 50μL of 100mM phosphate, pH 2.5 at 0.3°C. Immediately after the samples were quenched, 95μL of the sample was loaded onto an Acquity M-class UPLC (Waters) with sequential inline pepsin digestion (Waters Enzymate BEH Pepsin column, 2.1mm × 30mm) for 3min at 15°C followed by reverse phase purification (Acquity UPLC BEH C18 1.7μm at 0.2°C). Sample was loaded onto the column equilibrated with 95% water, 5% acetonitrile, and 0.1% formic acid at a flow rate of 40μL/min. A 7min linear gradient (5%–35% acetonitrile) followed by a ramp and 2min block (85% acetonitrile) was used for separation and directly continuously infused onto a Synapt XS using Ion Mobility (Waters). [Glu1]-Fibrinopeptide B was used as a reference.

Data from nondeuterated samples were used for peptide identification with ProteinLynx Global Server 3.0 (Waters). Full coverage of the protein was obtained, with the exception of the region from residues 289-296, where peptides were not detected. The filtered peptide list and MS data were imported into HDExaminer (Sierra Analytics) for deuterium uptake calculation using both retention time and mobility matching. Representative peptides were utilized for a final cumulative sequence coverage of 91.4%. Normalized deuterium uptake data was calculated for protein alone and with G10, and differential protection, defined as those regions with an average of 5% difference in deuteration between states over the 150-15000s timepoints, were mapped onto the crystal structure using PyMOL (Schrodinger).

### Native Mass Spectrometry (MS)

Stock solutions of BIg-*Rb*CBM74-BIg and Sas6 were de-salted and solvent exchanged into 200mM ammonium acetate (pH 6.8 – 7.0) using Amicon Ultra-0.5mL centrifugal filters (MilliporeSigma) with a 10kDa molecular weight cut-off. Ten consecutive washing steps were performed to achieve sufficient desalting. The final concentrations of each protein stock solution after desalting were estimated via UV absorbance at 280nm. A stock solution of G10 was prepared by dissolving a known mass in 200mM ammonium acetate to achieve a final concentration of 200μM. For native MS titration experiments used to quantify Kd values, the concentration of protein was fixed at 5μM, and enough G10 was added to achieve final concentrations of 0, 5, 25, 50, 100, and 150μM. Protein-G10 mixtures were then incubated at 4°C overnight to achieve equilibration prior to native MS analysis.

All native binding experiments were performed using a Q Exactive Orbitrap MS with Ultra High Mass Range (UHMR) platform (Thermo Fisher Scientific) [80]. Each sample (∼3µM) was transferred to a gold-coated borosilicate capillary needle (prepared in house), and ions were generated via direct infusion using a nano-electrospray ionization (nESI) source operated in positive mode. The capillary voltage was held at 1.2kV, the inlet capillary was heated to 250°C, and the S-lens RF level was kept at 80. Low m/z detector optimization and high m/z transfer optics were used, and the trapping gas pressure was set to 2. In-source trapping was enabled with the desolvation voltage fixed at -25V for improved ion transmission and efficient salt adduct removal. Transient times were set at 128ms (resolution of 25,000 at m/z 400), and 5 microscans were combined into a single scan. A total of ∼50 scans were averaged to produce the presented mass spectra. All full scan data were acquired using a noise threshold of 0 to avoid pre-processing of mass spectra. A total of three measurements for each ligand concentration were performed. Data were then processed and deconvoluted using UniDec software [81].

### K_d_ Measurements by Native MS

We performed titration experiments for both BIg-*Rb*CBM74-BIg and Sas6T using G10 and acquired modeled titration curves. Each bound state differed by ∼1639 Da, which agrees with the theoretical mass of G10. To obtain the binding constants, we summed the peak intensities of all abundant charge states in our mass spectra. *K*d values were calculated using the relative intensities of unbound protein and each ligand bound species from the mass spectra as previously described [82]. Briefly, the protein-ligand binding equilibrium of BIg-*Rb*CBM74-BIg with G10 in solution can be described by the following reversible reaction:

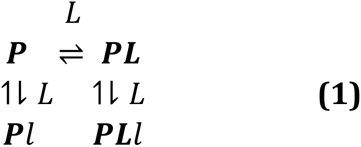

where *𝑳* is the ligand and *𝑷* and *𝑷𝑳* are the free protein and protein with one specifically bound ligand, respectively. BIg-*Rb*CBM74-BIg possesses one ligand-binding site, *Rb*CBM74. As the concentration of ligand is increased, ligand molecules can bind nonspecifically during the nESI process, generating artifactual peaks in the mass spectra corresponding to a two ligand-bound complex. As the concentration of ligand is increased, ligand molecules can bind nonspecifically during the nESI process, generating artifactual peaks in the mass spectra corresponding to a two ligand-bound complex. Here, we presume that nonspecific binding arises equally for free protein and that which possesses one specifically bound ligand represented by *𝑷𝑙* and *𝑷𝑳𝑙* in Eq. 1. Based on these assumptions, the equations of mass balance and binding states can be described the following system of equations:

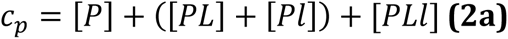

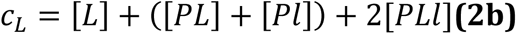

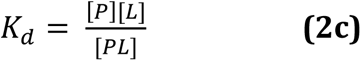

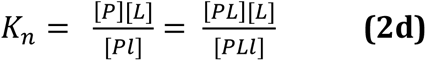

where *c_P_* and *c_L_* represent the total concentrations of protein and ligand, respectively, and concentrations in brackets represent those at equilibrium. *𝐾d* and *𝐾n* represent the dissociation constants for specific and nonspecific binding steps, respectively. If 1) the peak intensities of free protein and ligand-bound complexes are proportional to the abundances of those in solution and 2) the spray and detection efficiency of all species is the same, then the fractional intensities of each species can be determined:

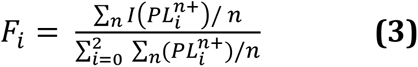

Here, the fractional intensities are calculated as the sum of the intensities of main peak ions at all charge states. Since a Fourier transform MS method is utilized, signal intensities are proportional to both ion abundance and charge state. Therefore, the ion intensities are normalized for each charge state, *n* [83, 84]. These fractional intensities can be calculated from the titration experiment at each ligand concentration and can then be related to the equilibrium constants:

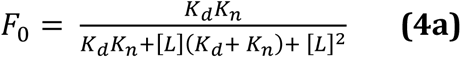

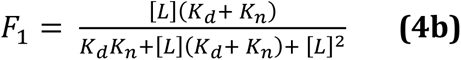

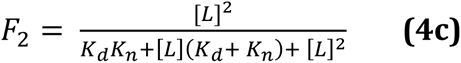

[L] can also be determined from nESI-MS titration data:

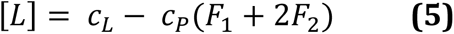

[L] was then obtained at each ligand concentration and applied to the Eqs. 4a-c. Equations 4a-b were then fitted to experimental fractional intensities using nonlinear least-squares curve fitting using the *lsqnonlin.m.* function in MATLAB. A more detailed derivation of these equations is provided elsewhere [82], along with the approach utilized for Sas6 which possesses two sites for specific binding (*Rb*CBM74 and *Rb*CBM26) and exhibits a third nonspecific bound state as shown in Eq. 6.

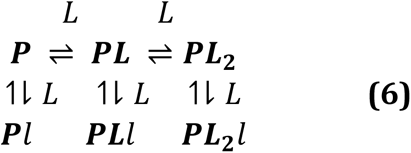

### Sequence collection

Amino acid sequences of CBM74 modules were collected according to information in the CAZy database (http://www.cazy.org/) yielding 29 sequences (CAZy update: March 2022) [85]. This set was subsequently completed with sequences of hypothetical CBM74s based on protein BLAST searches (https://blast.ncbi.nlm.nih.gov/Blast.cgi) using the CBM74 sequences from Sas6 of *Ruminococcus bromii* (GenBank Acc. No.: PKD32096.1) and the GH13_32 α-amylase from *Microbacterium aurum* (GenBank Acc. No.: AKG25402.1) as queries [20, 86, 87]. In total, three searches with each query sequence were performed, limiting the searched databases to taxonomy kingdoms of Bacteria, Archaea and Eucarya (with no relevant results for the latter two). To capture a wide spectrum of organisms harboring a CBM74 module, one non-redundant amino acid sequence was selected to represent each species and/or bacterial strain. The BLAST searches thus yielded 93 additional CBM74 sequences of bacterial origin; the last sequence taken being the CBM74 module of a putative α-amylase from uncultured *Eubacterium* sp. (GenBank: SCJ65691.1; E-value: 3e-39). That preliminary set of 122 sequences was reduced by eliminating 23 sequences due to their redundancy and/or incompleteness of the CBM74 module. The final set of CBM74 modules consisted of 99 sequences (**Extended Data Table 2**). All sequences were retrieved from the GenBank (https://www.ncbi.nlm.nih.gov/genbank/) and/or UniProt (https://www.uniprot.org/) databases [88, 89].

### Sequence comparison and evolutionary analysis

The alignment of 99 CBM74 modules from the final set was performed using the program Clustal-Omega (https://www.ebi.ac.uk/Tools/msa/clustalo/) [90]. Only a subtle manual tuning of the computer-produced alignment was necessary to perform to maximize sequence similarities. The evolutionary tree of these 99 sequences was calculated by a maximum-likelihood method (on the final alignment including the gaps) using the WAG substitution model and the bootstrapping procedure with 500 bootstrap trials implemented in the MEGA-X package [91–93]. The calculated tree file was displayed with the program iTOL [94] (https://itol.embl.de/). From both the alignment and the tree of all 99 sequences, a sample of 33 representative CBM74s was selected for a simplified alignment and tree. The structural comparison was created using the above-mentioned alignment in conjunction with the web-based CONSURF tool [54–56].

### Data availability

The X-ray structures and diffraction data reported in this paper have been deposited in the Protein Data Bank under the accession codes 7UWU, 7UWV and 7UWW. The SAXS data are deposited in the small angle x-ray scattering database (SASDB) under the accession code SASDPE2 [95]. All mass spectrometry data will be made available upon request.

## Funding and acknowledgements

This work is primarily supported by a Ruth L. Kirschstein National Research Service Award Individual Predoctoral Fellowship (F31 – F31AT011282 to A.L.P.) from the National Center for Complementary & Integrative Health (NCCIH) and a Research Program Project grant (P01-HL149633 to N.M.K.) from the National Heart, Lung, and Blood Institute (NHLBI) of the National Institutes of Health (NIH). Next-generation Native Mass Spectrometry technologies were supported by National Institute of General Medical Sciences (NIGMS) of the NIH (R01-GM095832 to B.T.R.). Hydrogen-Deuterium exchange mass spectrometry acquisition was supported by National Science Foundation (NSF) (DBI 2018007 to C.W.V.K.). The structural biology approaches herein used resources of the Advanced Photon Source; a U.S. Department of Energy (DOE) Office of Science User Facility operated for the DOE Office of Science by Argonne National Laboratory under Contract No. DE-AC02-06CH11357. The Biophysics Collaborative Access Team is supported by P30-GM138395 from NIGMS-NIH. Use of the Pilatus 3 1M detector was provided by Grant 1S10OD018090-01 from NIGMS-NIH. Use of the LS-CAT Sector 21 was supported by the Michigan Economic Development Corporation and the Michigan Technology Tri-Corridor (Grant 085P1000817). S.J. and F.M. thank the Slovak Grant Agency VEGA for the financial support by the Grant No. 2/0146/21. In collaboration with this research, we acknowledge support from the University of Michigan Biomedical Research Core Facilities Light Microscopy Core. For the native mass spectrometry work, we would like to acknowledge the Biological Mass Spectrometry facility at the University of Michigan Department of Chemistry. The content is solely the responsibility of the authors and does not necessarily represent the official views of VEGA, the National Science Foundation, or National Institutes of Health.

## Author contributions

N.M.K. and A.L.P. conceptualization;

A.L.P., F.M.C., R. V-V., K.M.A., F.M., T.C., Z.W., J.H., C.W.V.K, S.J., B.T.R., and N.M.K. data curation;

A.L.P., F.M.C., R.V-V., K.M.A., F.M., T.C., Z.W., J.H., C.W.V.K, S.J., B.T.R., and N.M.K. formal analysis and data interpretation;

N.M.K., A.L.P., C.W.V.K, S.J., B.T.R., Z.W., and J.H. funding acquisition;

A.L.P., F.M.C., R. V-V., K.M.A., F.M., T.C., Z.W., J.H., C.W.V.K, and S.J. investigation;

A.L.P., F.M.C., R. V-V., K.M.A., F.M., T.C., Z.W., J.H., C.W.V.K, S.J., B.T.R., and N.M.K. methodology;

A.L.P., F.M.C., R.V-V., C.W.V.K, S.J. and N.M.K original draft;

A.L.P., F.M.C., R.V-V., F.M., J.H., C.W.V.K, S.J., B.T.R., and N.M.K. writing-review and editing;

N.M.K., J.H., C.W.V.K, S.J., and B.T.R. supervision;

A.L.P., F.M.C., R.V-V., T.C., and S.J. visualization.

## Extended Data

**Extended Table 1A:** Average masses assigned to native mass spectrometry peaks

| P | 0 bound (Da) | 1 bound (Da) | 2 bound (Da) | 3 bound (Da) | Av Diff. (Da)<sup>a</sup> |
| --- | --- | --- | --- | --- | --- |
| Blg-RbCBM74-Big | 57635.3 ± 0.8 | 59274.7 ± 0.6 | 60913.8 ± 0.2 | N/A | 1639.4 ± 0.8 |
| Sas6 | 69064.9 ± 0.6 | 70704.3 ± 0.5 | 72343.8 ± 1.8 | 73982.1 ± 0.6 | 1639.3 ± 1.0 |

<sup>a</sup>Average difference between bound states across all ligand concentrations.

**Extended Table 1B:** Binding parameters determined by Native Mass Spectrometry

| P | Charge States | K<sub>d1</sub> (μM) | K<sub>d2</sub> (μM) | K<sub>n</sub> (μM)<sup>a</sup> | SSR<sup>b</sup> |
| --- | --- | --- | --- | --- | --- |
| Blg-RbCBM74-Big | 11+ to 14+ | 3.8 ± 0.5 | N/A | 1154.4 ± 378.0 | 0.0111 |
| Sas6 | 13+ to 15+ | 3.4 ± 0.5 | 165.6 ± 38.8 | 782.6 ± 647.9 | 0.0170 |

<sup>a</sup>K<sub>n</sub> - dissociation constant for nonspecific binding step during nESI. Values reported with 95% confidence interval.
<sup>b</sup>SSR - sum of squared residuals

**Extended Table 2:**
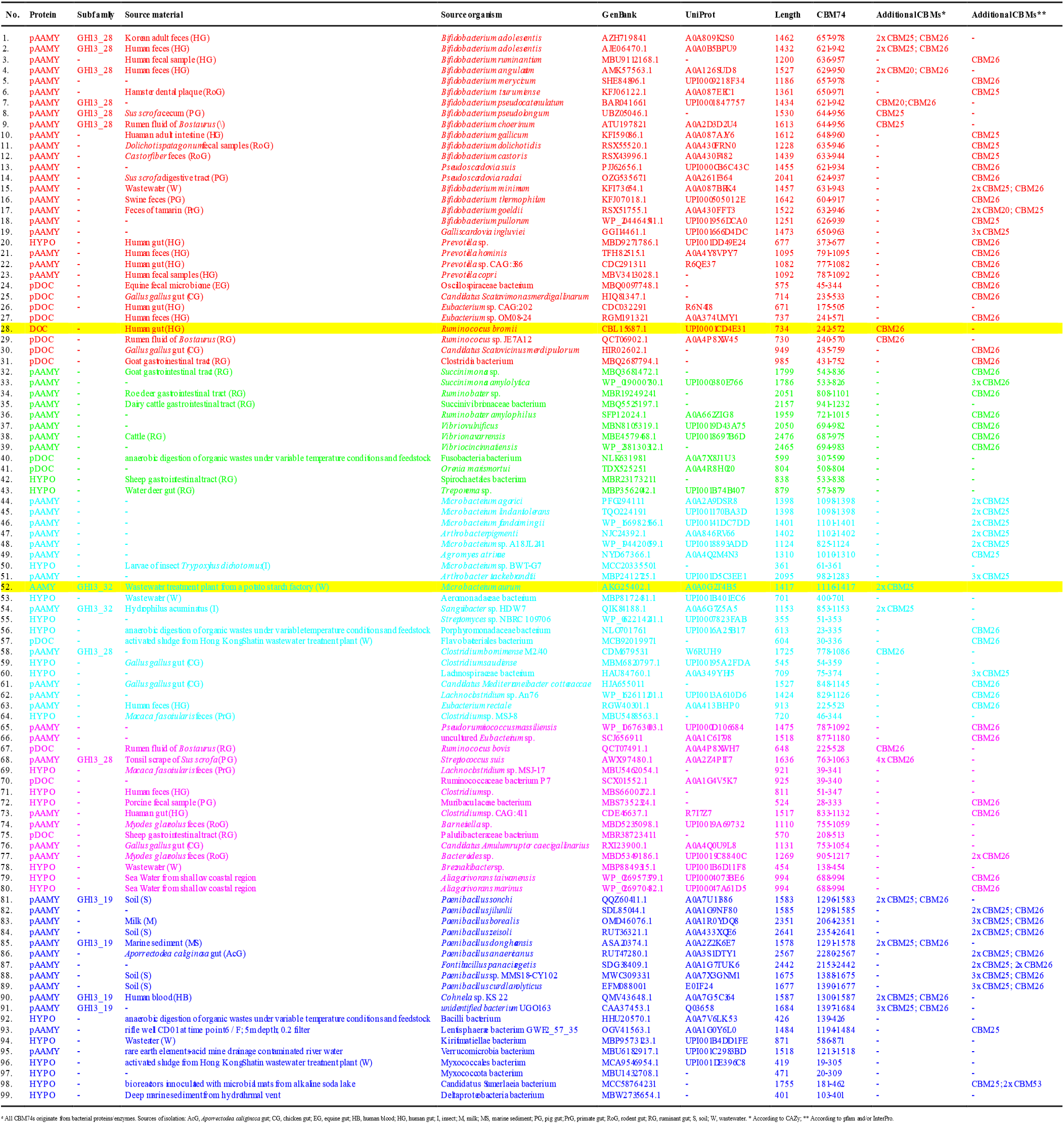
List of 99 selecte d CBM74 sequences.*^a^*

**Extended Data Figure 1:**
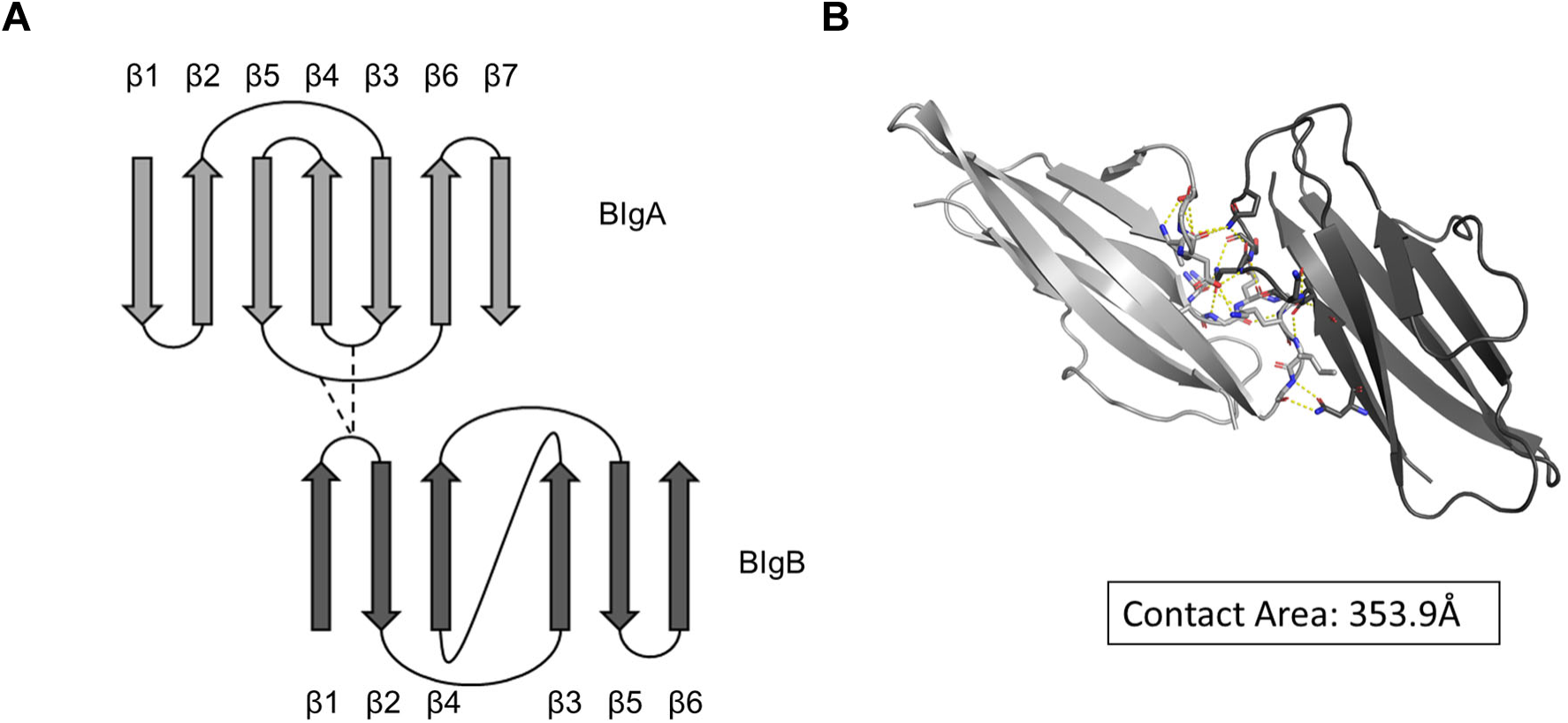
Bacterial Ig-like domains of Sas6 interact via extensive hydrogen bonding. **A.** Topology map of BIgA and BIgB domains illustrating the Greek key motif in BIgA and showing the loops that hydrogen bond with one another. **B.** A surface area analysis of the BIg domains using PISA in CCP4 gives a buried surface area of 353.9Å [37]. Residues providing hydrogen bonding are represented by stick side chains and the hydrogen bonds are shown by dashed yellow lines.

**Extended Data Figure 2:**
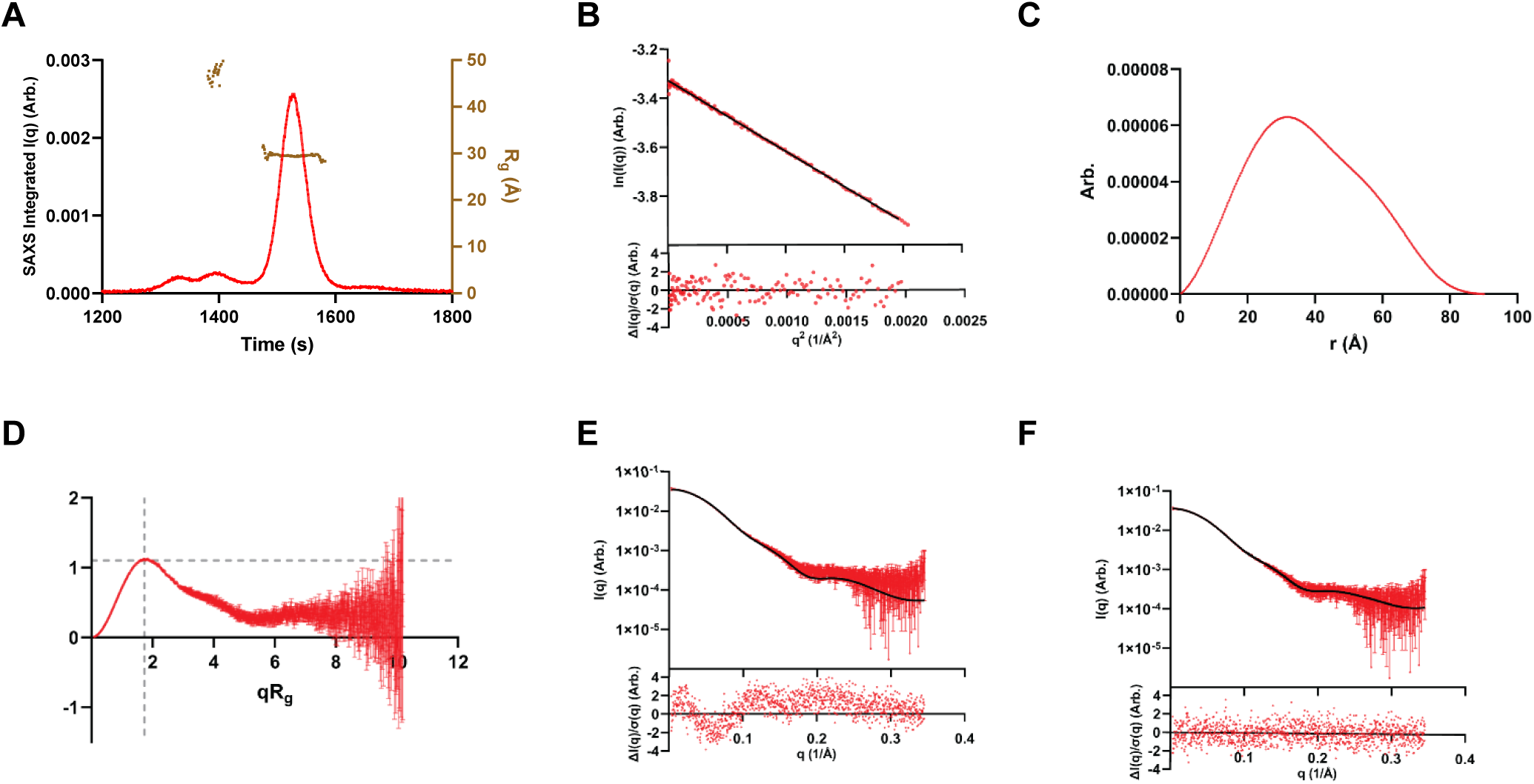
Small Angle X-Ray Scattering indicates that Sas6 remains mostly compact in solution with minor extension beyond that of the crystal structure. **A.** Total subtracted scattering intensity (left y axis) and Rg (right y axis) as a function of time for the SEC-SAXS elution. **B.** Guinier fit analysis with normalized residual shown in the bottom panel. **C.** P(r) versus r normalized by I(0). **D.** Dimensionless Kratky plot; y=3/*e* and x=*√*3 as dashed gray lines to indicate where a globular protein would peak. **E.** FoXS and **F.** MultiFoXS fits (black) to the Sas6T SAXS data (red) with normalized residual shown in the bottom panel.

**Extended Data Figure 3:**
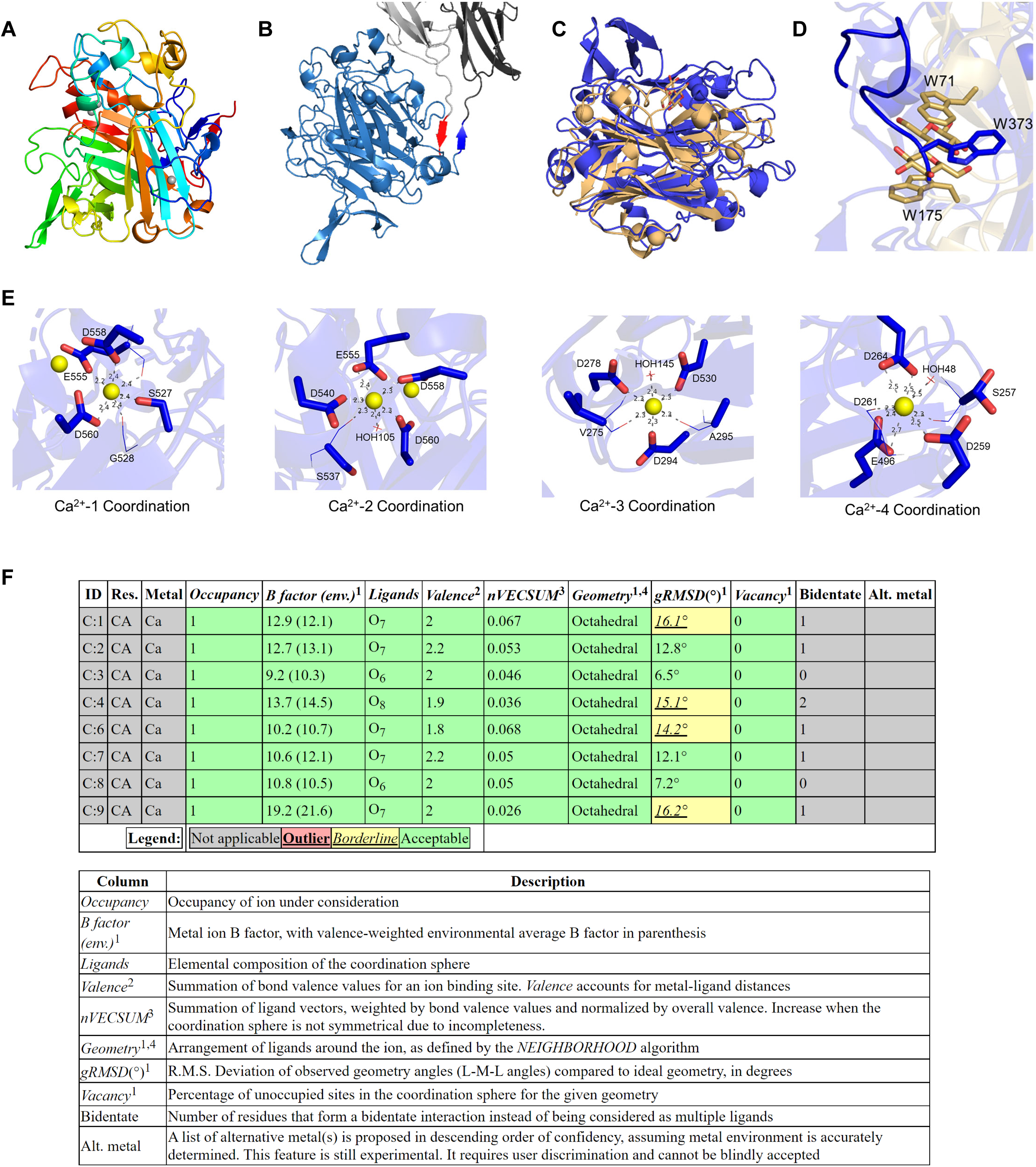
*Rb*CBM74 is a singular globular domain, most similar to *Tm*CBM9. **A.** Structure of *Rb*CBM74 (PDB *7uww*) colored from N-terminus (blue) to C-terminus (red). **B.** Short β-strands leading into and out of *Rb*CBM74 domain are colored in red and blue. **C.** Overlay of *Tm*CBM9 (gold) (PDB *1i82*-A) and *Rb*CBM74 (blue). The DALI server calculated an RMSD of 3.2Å and sequence identity of 17%. **D.** Close-up view of *Tm*CBM9 binding site showing the two *Tm*CBM9 Trp residues involved in binding cellobiose (gold) and W373 of *Rb*CBM74 (blue) which lies in the same region but is occluded from the surface by a loop containing residues 374-384. **E.** Zoomed in view of calciums coordinated in the *Rb*CBM74 domain with side chains shown in sticks, main chain shown in lines and Ca^2+^ ions by yellow spheres. Atomic distances are shown in Å and residues are labeled. Residues are colored by element with oxygen shown in red. **F.** Ion validation by web server CheckMyMetal [72].

**Extended Data Figure 4:**
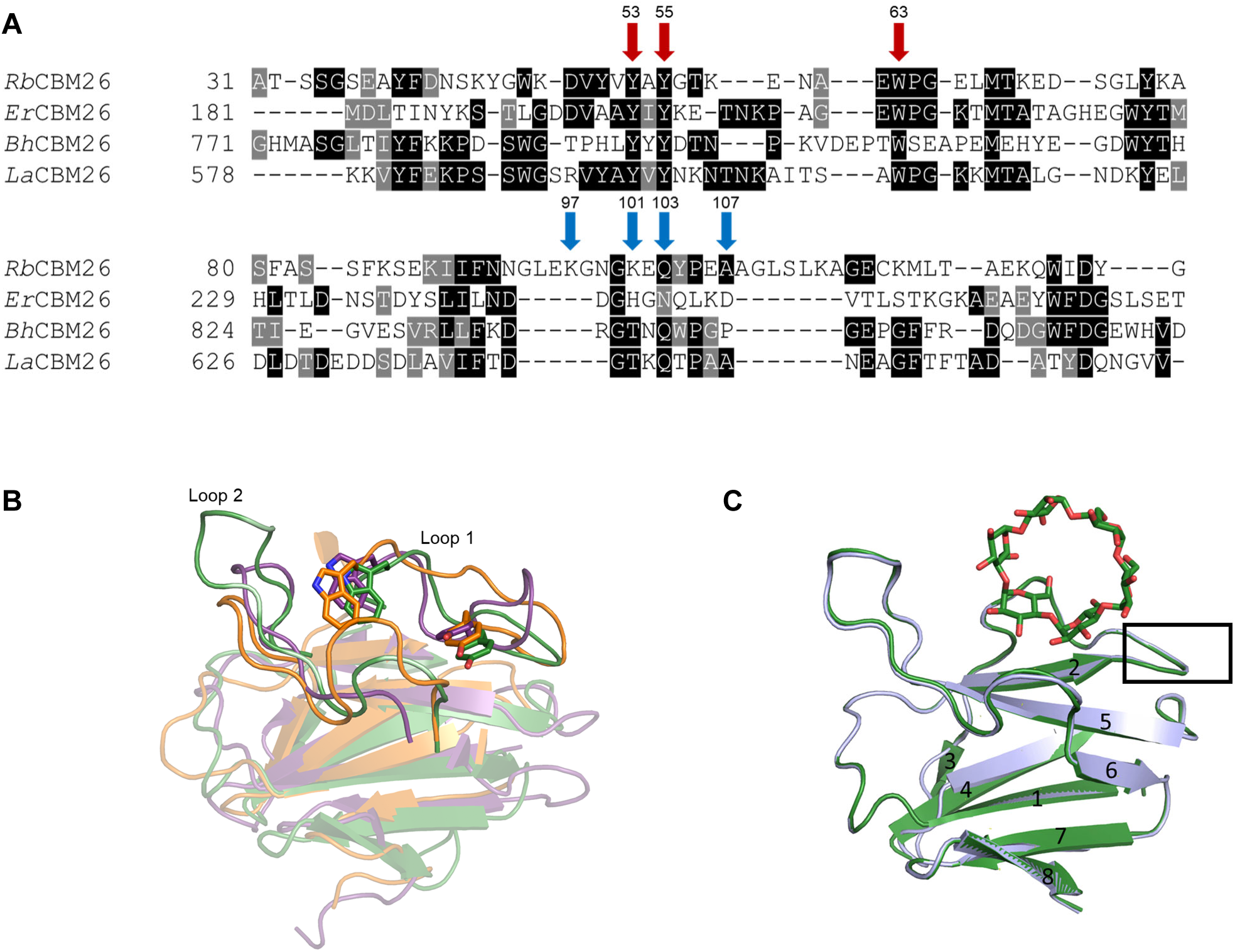
RbCBM26 shares a conserved binding site with other CBM26 family members. **A.** Sequence alignment of RbCBM26 (RBL236_00020), *Er*CBM26 (ERE_20420), *Bh*CBM26 (BH0413), and *La*CBM26 (Q48502). Conserved binding site residues are indicated by a red arrow while variable residues are indicated by a blue arrow and provide hydrogen bonding. **B**. Overlay of *Rb*CBM26 (green) with *Bacillus halodurans* CBM26 (PDB *2c3h*, orange), and *Eubacterium rectale* Amy13K CBM26 (PDB *6b3p*, purple). **C**. Overlay of unliganded *Rb*CBM26 (blue) and ACX-bound *Rb*CBM26 (green) showing that loop 1 does not move upon ligand binding. β-strands are numbered for reference.

**Extended Data Figure 5:**
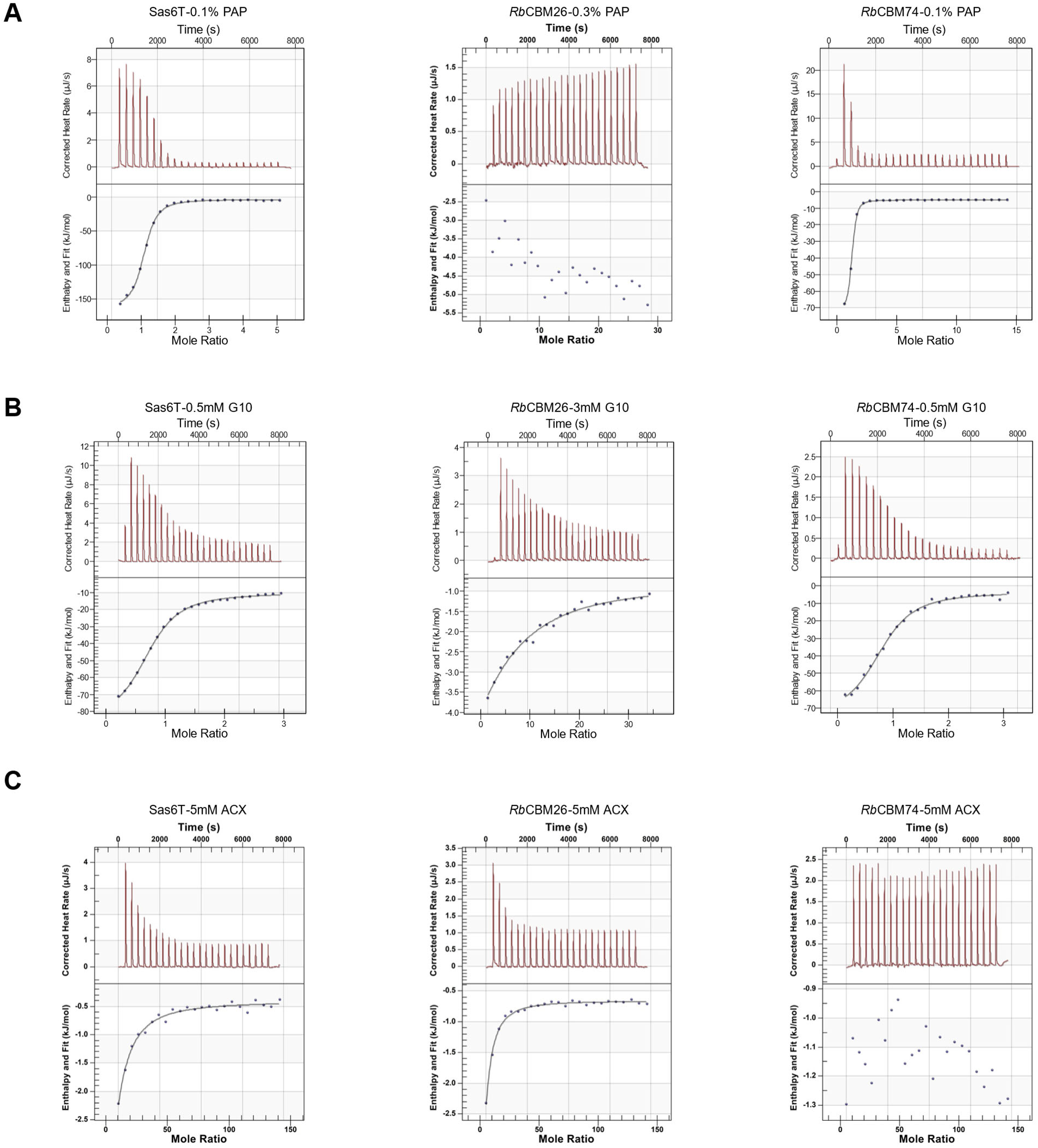
Representative ITC graphs of Sas6 domains. Sas6T, *Rb*CBM74, and BIg-*Rb*CBM74-BIg binding to **A.** potato amylopectin, **B.** maltodecaose (G10), and C. α-cyclodextrin (ACX).

**Extended Data Figure 6:**
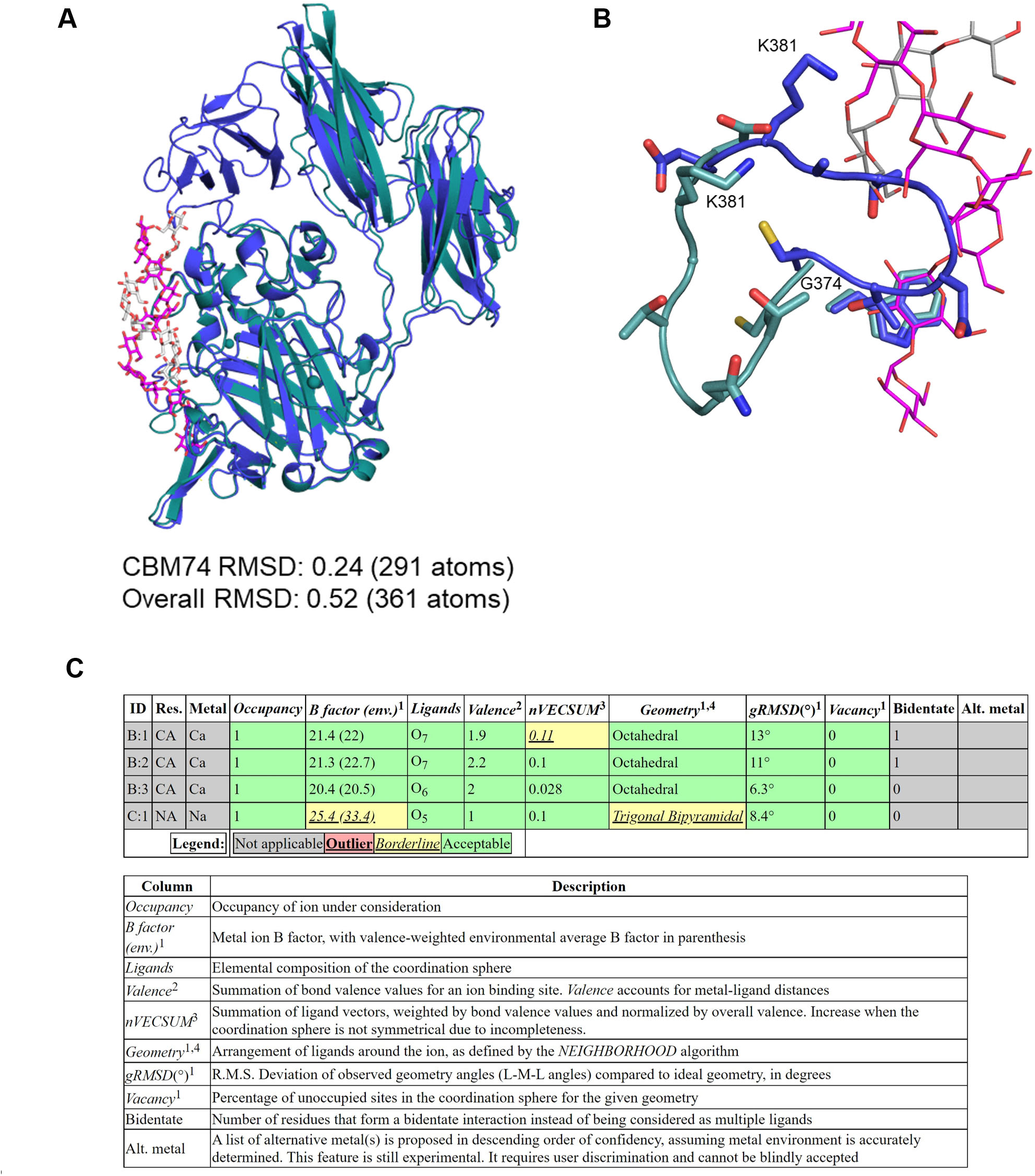
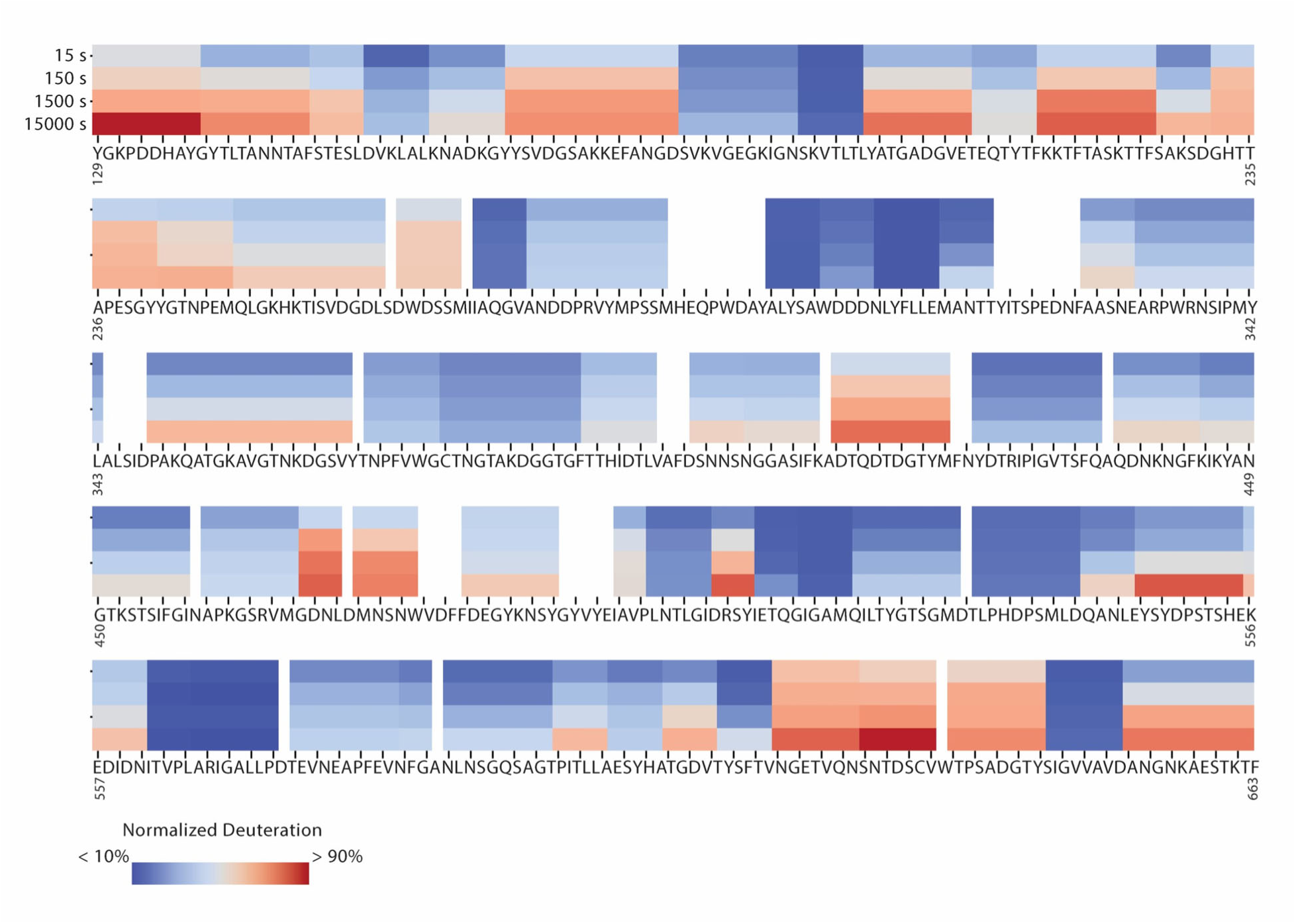
*Rb*CBM74 undergoes minor conformational changes upon ligand binding. **A.** Overlay of *Rb*CBM74 from Sas6T structure (PDB *7uww*) in blue with *Rb*CBM74 from BIg-*Rb*CBM74-BIg co-crystal structure (PDB *7uwv*) in deep teal. **B.** Loop from G374-G382 demonstrating that the unliganded loop (blue) occludes W373 but moves to allow access to W373 in the ligand-bound structure (deep teal). **C.** Validation of ion identities with CheckMyMetal [72]. Note Ca^2+^-4 is exchanged for a Na^+^ ion in the G10 liganded structure.

**Extended Data Figure 7:**
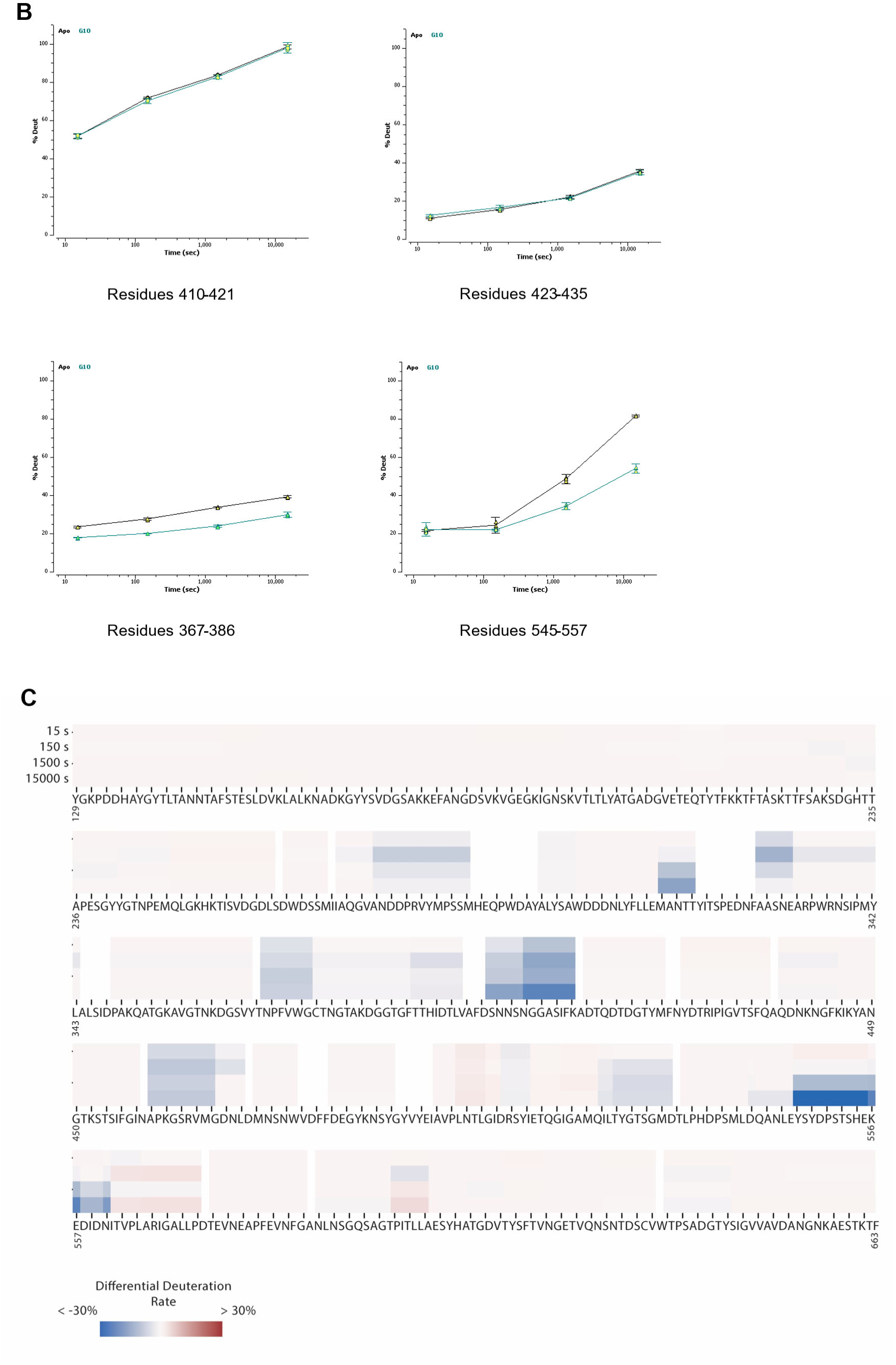
HDX-MS analysis of *Rb*CBM74. **A.** Heatmap of exchange dynamics of BIg-*Rb*CBM74-BIg. All values are the average of three replicates. **B.** Representative differential uptake for peptides that both showed no significant difference (upper panels) and those which showed significant differential decreased deuteration (lower panels) in the G10 bound BIg-RbCBM74-BIg. Data points are represented by the mean +/-standard deviation. **C.** Heatmap of the differential exchange dynamics of BIg-*Rb*CBM74-BIg in the absence and presence of G10. Blue represents lower exchange (protection) in the G10 bound form and red higher exchange in the G10 bound form. All values are the average of three replicates.

**Extended Data Figure 8:**
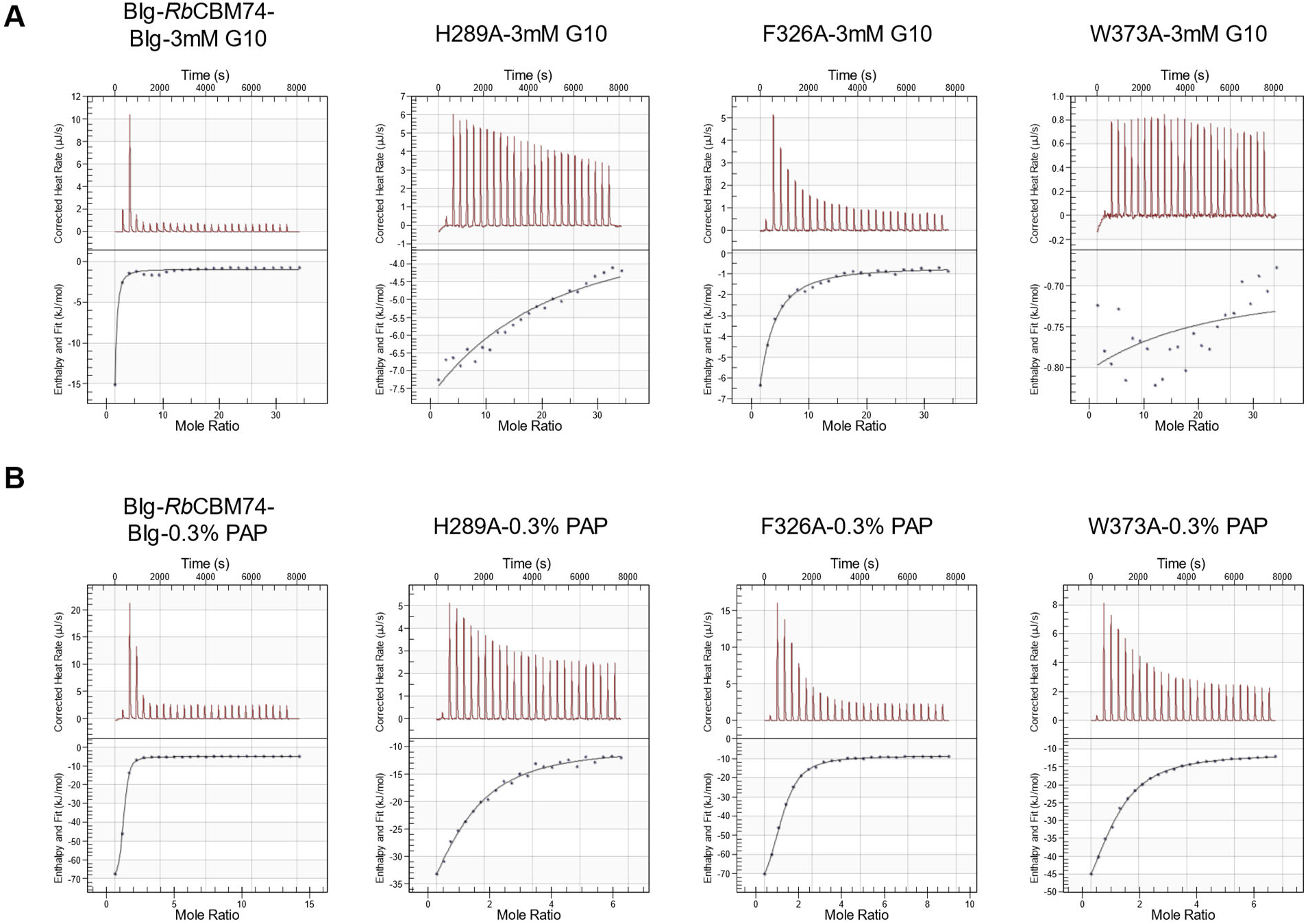
Representative ITC graphs of *Rb*CBM74 mutations. BIg-*Rb*CBM74-BIg, H289A, F236A, and W373A mutations binding to **A.** maltodecaose (G10), and **B.** potato amylopectin (PAP).

**Extended Data Figure 9:**
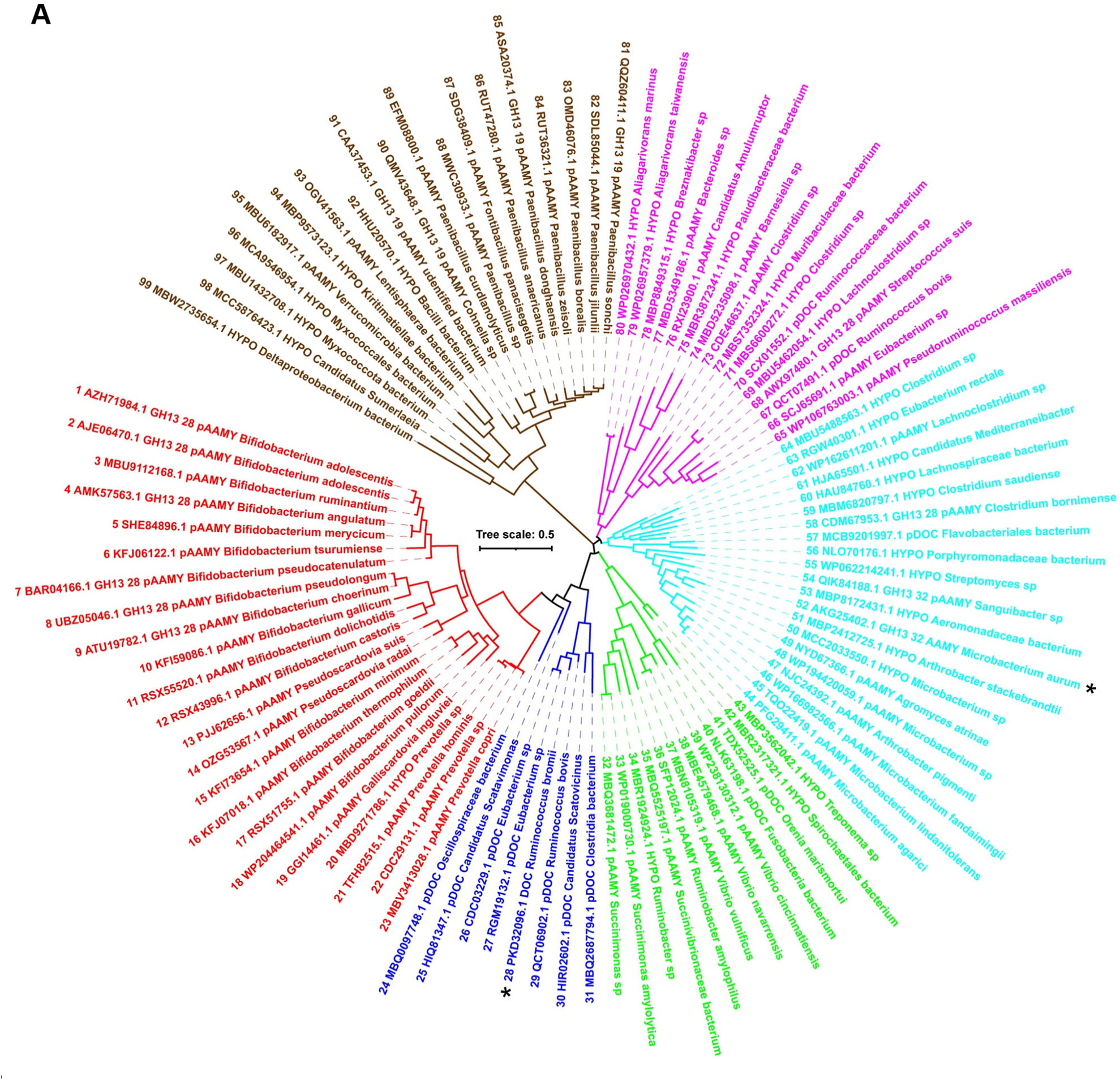
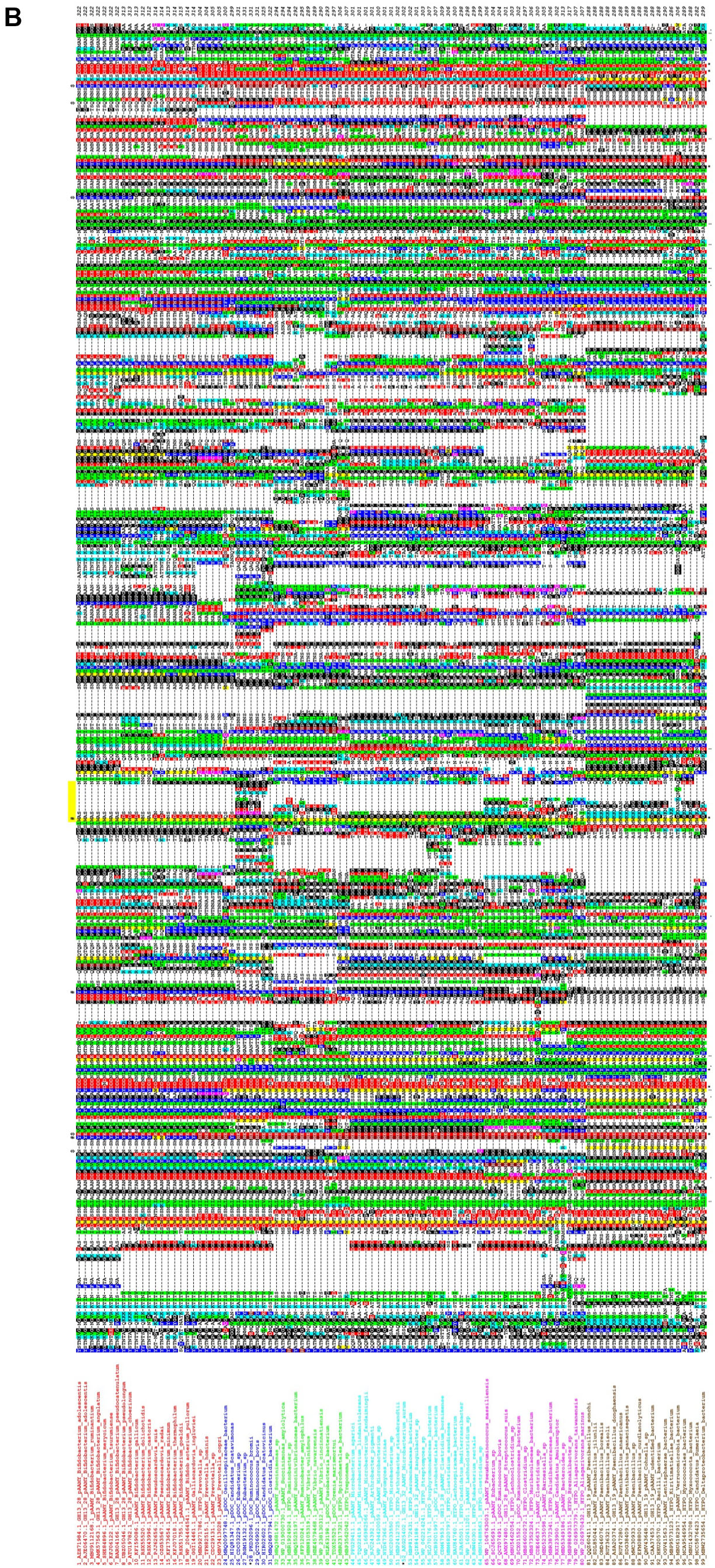
Conservation of binding residues among all 99 CBM74 family members. **A.** A maximum-likelihood tree covering 99 sequences with emphasis on the two experimentally characterized CBM74s, Sas6 from *Ruminococcus bromii* (No. 28, blue cluster) and the subfamily GH13_32 α-amylase from *Microbacterium aurum* (No. 52; cyan cluster). For details concerning all 99 CBM74 sequences, see Extended Data Table 2. A simplified tree showing 33 selected CBM74 sequences representing all clusters is shown in Fig. 6A. **B.** Sequence alignment of the 99 CBM74 sequences. The labels of protein sources consist of the order number (1-99), GenBank accession number, abbreviation of the source protein/enzyme and the name of the organism. The two experimentally characterized CBM74 are marked by an asterisk. The six individual groups distinguished from each other by different colors correspond to six clusters seen in the evolutionary tree (panel A); the sequence order in the alignment (starting from the top from 1 to 99) reflects their order in the tree in the anticlockwise manner (starting from the first sequence in the red cluster). The residues responsible for stacking interactions and involved in hydrogen bonding with glucose moieties of the bound α-glucan are signified by a hashtag and a dollar sign, respectively, above the alignment. The flexible loop observed in the three-dimensional structure of *Rb*CBM74 is highlighted by the short yellow strip over the alignment. Identical and similar positions are signified by asterisks and dots/semicolons under the alignment blocks. The color code for the selected residues: W, yellow; F, Y – blue; V, L, I – green; D, E – red; R, K – cyan; H – brown; C – magenta; G, P – black.

