## Supplementary figures and images for "The *Ruminococcus bromii* amylosome protein Sas6 binds single and double helical α-glucan structures in starch"

### Supplemental Tables

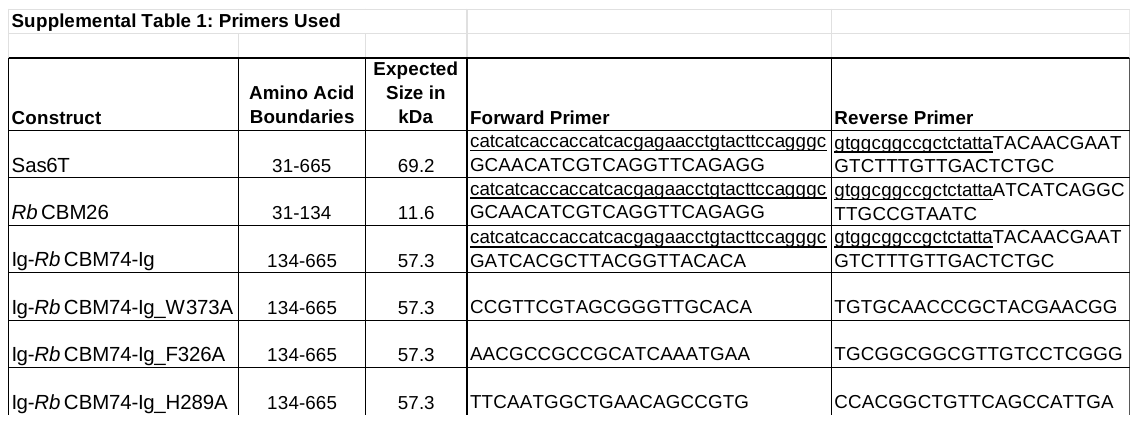

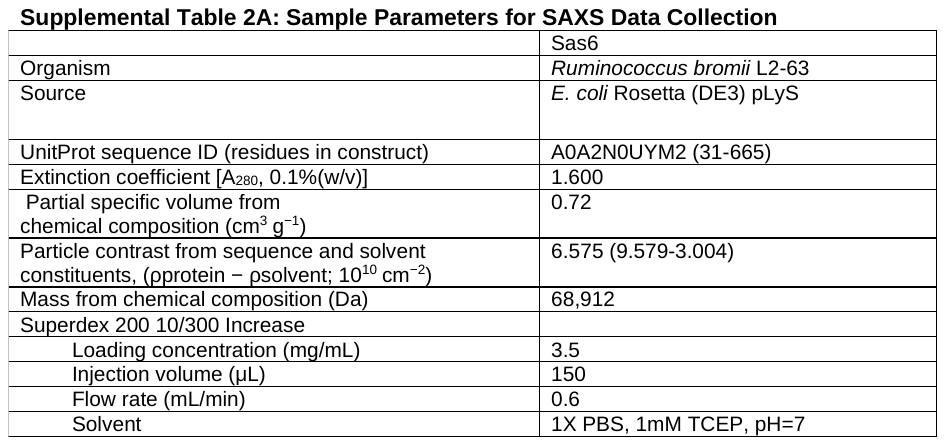

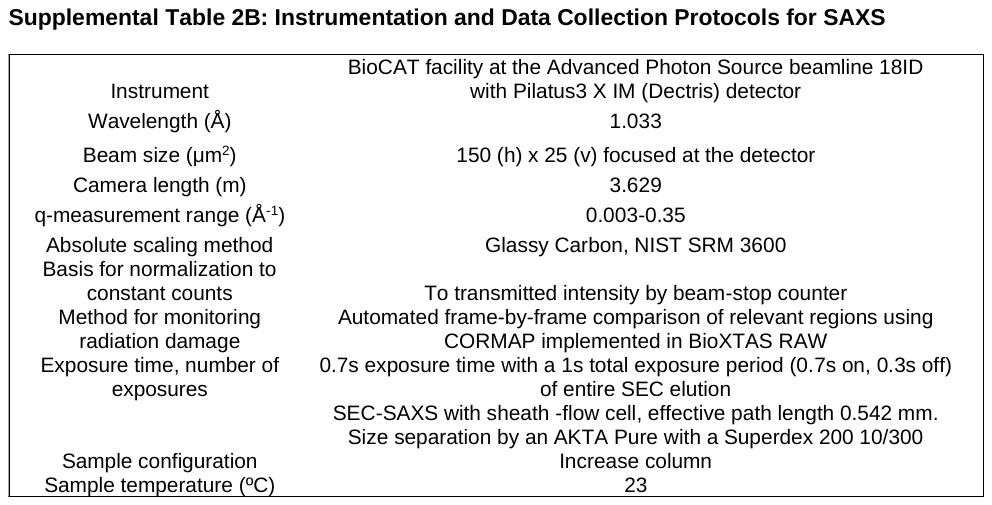

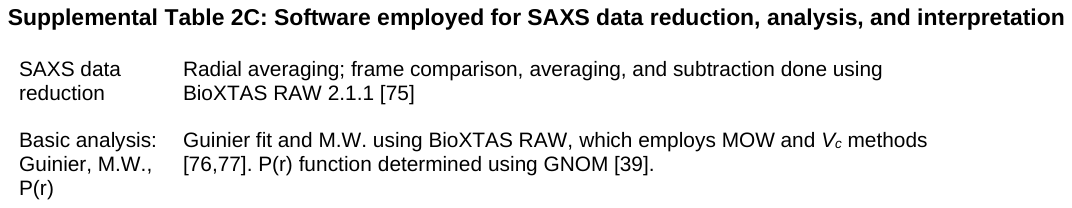
